# Chronic circadian disruption modulates breast cancer cell stemness and their immune microenvironment to drive metastasis in mice

**DOI:** 10.1101/777433

**Authors:** Eva Hadadi, William Taylor, Xiaomei Li, Yetki Aslan, Marthe Villote, Julie Rivière, Gaelle Duvallet, Charlotte Auriau, Sandrine Dulong, Isabelle Raymond Letron, Sylvain Provot, Annelise Bennaceur-Griscelli, Hervé Acloque

## Abstract

Breast cancer is the most common type and one of the major causes of cancer death in woman worldwide. Epidemiological studies have established a link between night shift work and increased cancer risk, suggesting that circadian disruption may interfere with carcinogenesis. We aim to shed light on the effect of chronic jetlag on mammary tumour development. Therefore, we used a mouse model of spontaneous mammary tumorigenesis that we exposed to chronic circadian disruption. We observed that circadian disruption significantly increases cancer cell dissemination and metastasis. It also enhances the stemness and tumour–initiating potential of tumour cells and creates an immunosuppressive shift in the tumour microenvironment. We finally showed that all these defects can be corrected by the use of a CXCR2 inhibitor. Altogether, our data provide a conceptual framework to better understand and manage the effects of chronic circadian disruption on breast cancer progression.

## Introduction

Breast cancer (BC) is the most frequent cancer in women worldwide. The cumulative risk for a woman to develop BC is around 5% worldwide and to die of it is 1.4%. There were more than 2 million newly diagnosed cases in 2018, representing almost 25% of all cancer cases among women. BC is the leading cause of death for women in most countries.^1^ While genetic causes account for less than 10% of BCs, non-hereditary causes are the major drivers of BCs development. This includes nutrition and alcohol consumption, exogenous hormone intake, reproduction and menstruation parameters.^1^ Environmental factors such as air pollution or altered light/dark cycles like for night shift work can also affect BC incidence.^2–4^ For the latter, the International Agency for Research on Cancer (IARC) classified circadian rhythm disruption (CRD) as probably carcinogenic in 2007 based on 8 epidemiologic studies and available data on animal models.^4^ Since then, additional and better-documented epidemiologic studies, genome-wide association studies (GWAS) and cellular and animal studies confirmed the link between BCs development and circadian disruption.^5^ A recent population-based case-control study confirms that factors including night work duration, length of shifts and time since last night shift affect the odd ratios for BC mostly in premenopausal women.^6^ The hormonal receptor status is also important and BC risk associated with night work is only higher for ER+ HER2+ cancer. These epidemiological studies are supported by GWAS studies showing a significant statistical association between genetic variations located in circadian genes (ARNTL, CLOCK, CRY1, CRY2, RORA, RORB, RORC, PER1) and the risk of breast cancer or between circadian clock genes expression and metastasis-free survival.^7,8^ Experimental studies on mammary epithelial cells also support an important role for circadian core clock genes on mammary gland formation and function. Per2^-/-^ mutant female mice fail to form normal terminal mammary ducts with an excess of basal progenitors.^9^ Arntl^-/-^ female mutants possess less ductal branches and ductal length and more terminal end buds while Clock^-/-^ mutants female present defect in daily maternal behaviour and milk production.^10,11^ Depletion of core circadian genes in human mammary epithelial cells confirm these *in vivo* observations and confirm a strong effect of these genes on the stemness of mammary epithelial cells, either by decreasing or increasing stemness.^9,12,13^ However, it is not always clear whether the observed phenotypes resulted from a non-circadian function of these transcription factors or an indirect consequence of a global dysregulation of the circadian clock in cells and tissues.

Experimental evidences on the effects of CRD on breast cancer initiation were previously described using an inducible p53 mutant mouse model prone to develop primary mammary tumours. While these mice develop mammary tumours in 50 weeks, onset was 8 weeks earlier when the mice experienced CRD and supports that CRD is a driving factor for breast cancer development.^14^

This last study only focused on the onset of tumorigenesis and we decided to explore the effects of CRD on tumour progression, cancer cell dissemination and immune phenotype. We used the MMTV:PyMT model of spontaneous murine mammary carcinogenesis ^15^ and tested the effects of a chronic CRD applied for 10 weeks at the beginning of the tumorigenesis at puberty.

## Results

### Chronic CRD moderately affects Primary tumour development and mice physiology

At 6 weeks old, mice carrying the MMTV:PyMT and MMTV:LUC transgenes were divided into 2 lots. One was maintained for 10 weeks in a normal alternation of light and dark periods (LD, 12h light and 12h dark) and one was exposed for 10 weeks to a chronic CRD through a continuous jet lag, corresponding to a shortening of 8 hours of each dark period out of two (JL) (Fig. 1A). This jet lag protocol was previously shown to totally disrupt the 24h-period of the rhythmic pattern of rest (12h Light period) and activity (12h Dark period) and is frequently used to mimick the effects of shift works or frequent eastbound transmeridian flights.^16^ From 10 weeks to the end of the experimentation at 16 weeks, growth of primary tumours was monitored every two weeks by imaging luciferase activity driven by the MMTV:LUC transgene. At 16 weeks, mice were weighted then sacrificed and tissues were processed for further analysis. The median weight of JL mice was slightly but not significantly increased compared to LD mice, as expected from previous studies (Fig. 1B).^17,18^ The peripheral blood cell counts were similar between the two experimental conditions (Fig. 1C). In addition, most of the blood biochemistry parameters (Supplementary Information, Fig. S1) showed no significant differences with the exception of glucose level, significantly decreased and total cholesterol, triglycerides and free fatty acids levels that are significantly increased in the plasma of these mice (Fig. 1D). *In vivo* imaging revealed no significant difference in the start of tumorigenesis (Fig. 1E). However, tumour growth showed slight increase with age and tumour burden was also moderately but significantly increased in JL mice (Fig. 1E and 1F). We assessed the grade of primary tumours on paraffin-embedded sections stained with HES (hematoxylin, eosin and saffron). Different tumour grades, ranging from hyperplasia to late carcinoma,^19,20^ were observed in primary tumours from LD and JL mice with a slight tendency to observe more malignant lesions in JL mice (Supplementary Information, Table S1 and Fig. S2).

**Figure 1:**
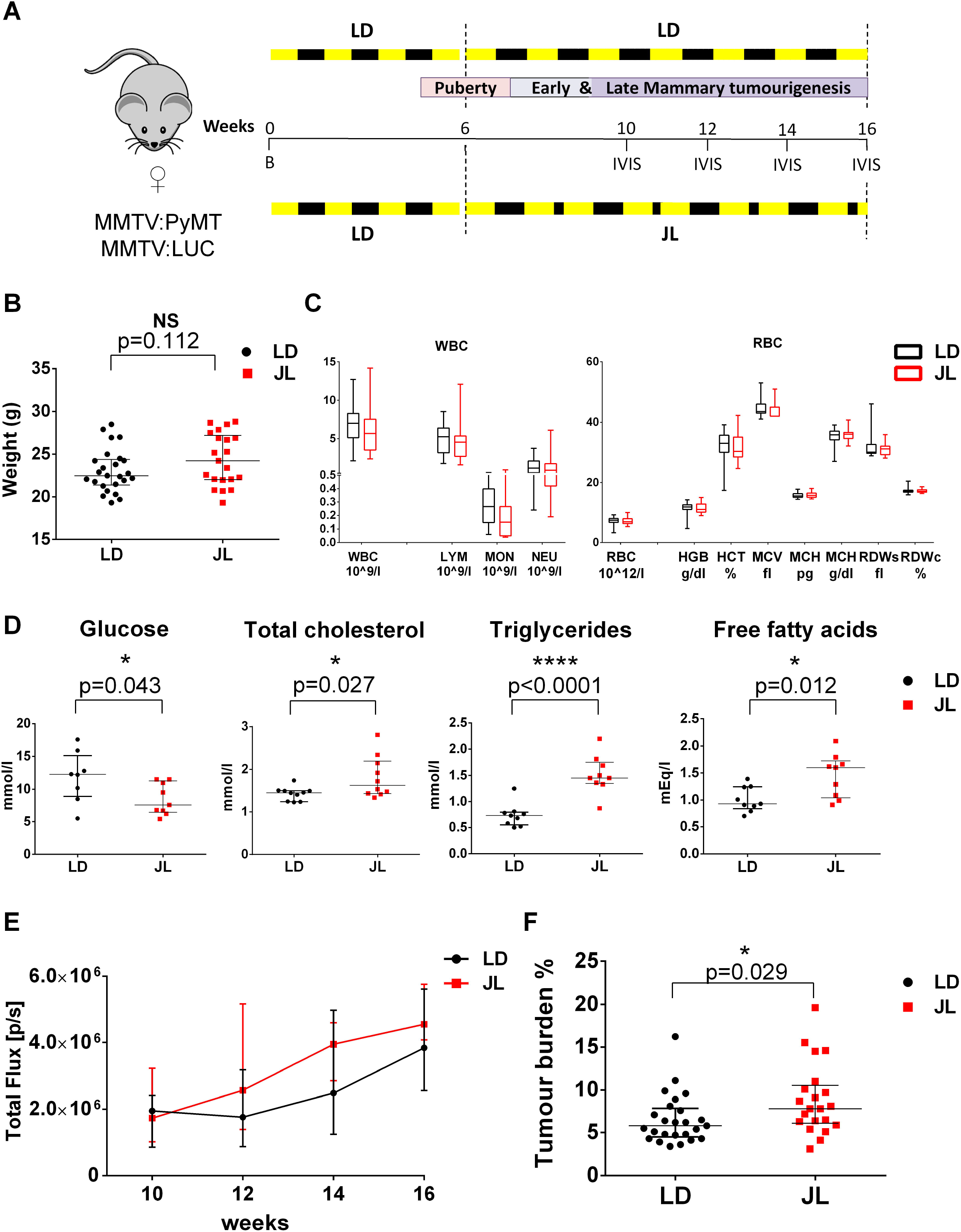
Chronic circadian disruption slightly enhances tumour burden. (A) Schematic graph of experimental timeline to assess the effect of chronic jet lag on spontaneous mammary tumorigenesis. LD: 12h light and 12h dark; JL: jet lag corresponding to shortening of 8 hours each dark period out of two; B: birth; IVIS: tumour growth monitoring using bioluminescence; S: sacrifice. Dash lines highlight the start and end points of the experimentation. (B) Weight of mice at sacrifice in LD (n=25) and JL (n=21). Data are presented as scatter dot plot with lines representing median with interquartile. p-value calculated from an unpaired t-test. (C) Blood cell counts: white blood cell (WBC) and red blood cell (RBC) total numbers in LD (n=16) and JL (n=17) groups. LYM: lymphocytes; MON: monocytes; NEU: neutrophils; HGB: haemoglobin; HCT: haematocrit; MCV: mean corpuscular volume; MCH: mean corpuscular haemoglobin; RDW: red cell distribution width. Data are presented as boxplots with whiskers representing min to max. (D) Glucose, total cholesterol, triglycerides and free fatty acids profile of LD and JL mice. Data are presented as scatter dot plot with lines representing median and interquartile. p-value calculated from an unpaired t-test. (E) Timelines of tumour growth in total flux [p/s] measured by *in vivo* bioluminescence imaging in LD (n=6) or JL (n=6) groups. (F) Tumour burden (tumour to body weight ratio) % in LD (n=25) or JL (n=21) conditions. Data are presented as scatter dot plot with lines representing median with interquartile. p-value calculated from an unpaired t-test.

### Chronic CRD promotes cancer cell dissemination and metastasis formation

We then explored if chronic CRD can affect the dissemination of cancer cells. We quantified the presence of disseminated cancer cells (DCCs) through the expression of the PyMT transgene by flow cytometry and real-time PCR. We observed a significant increase of DCCs in the bone marrow of JL mice by using both types of quantification with almost a two-fold increase of DCCs in BM Lin^-^ mono-nucleated cells (Fig. 2A). PyMT transgene expression was also higher in the blood stream of JL mice and flow analysis confirms an increase of circulating cancer cells (CTCs) (Fig. 2B). We also observed DCC like cells with H&E staining on bone sections while μCT analysis revealed abnormal bone lesions also highlighting the dissemination of cancer cells to bone (Supplementary Information, Fig. S3). In agreement with these observations, we also observed a significant increase of metastasis prevalence in JL mice compared to LD mice (Fig. 2C). The proportion of mice with lung metastasis goes through 28% in LD to 52% in JL (Fig. 2C) and the number of metastatic foci is also significantly increased in JL mice (Fig. 2D; Supplementary Information Fig. S3). Altogether, our results reveal a significant impact of chronic CRD on cancer cell dissemination and their metastatic potential.

**Figure 2:**
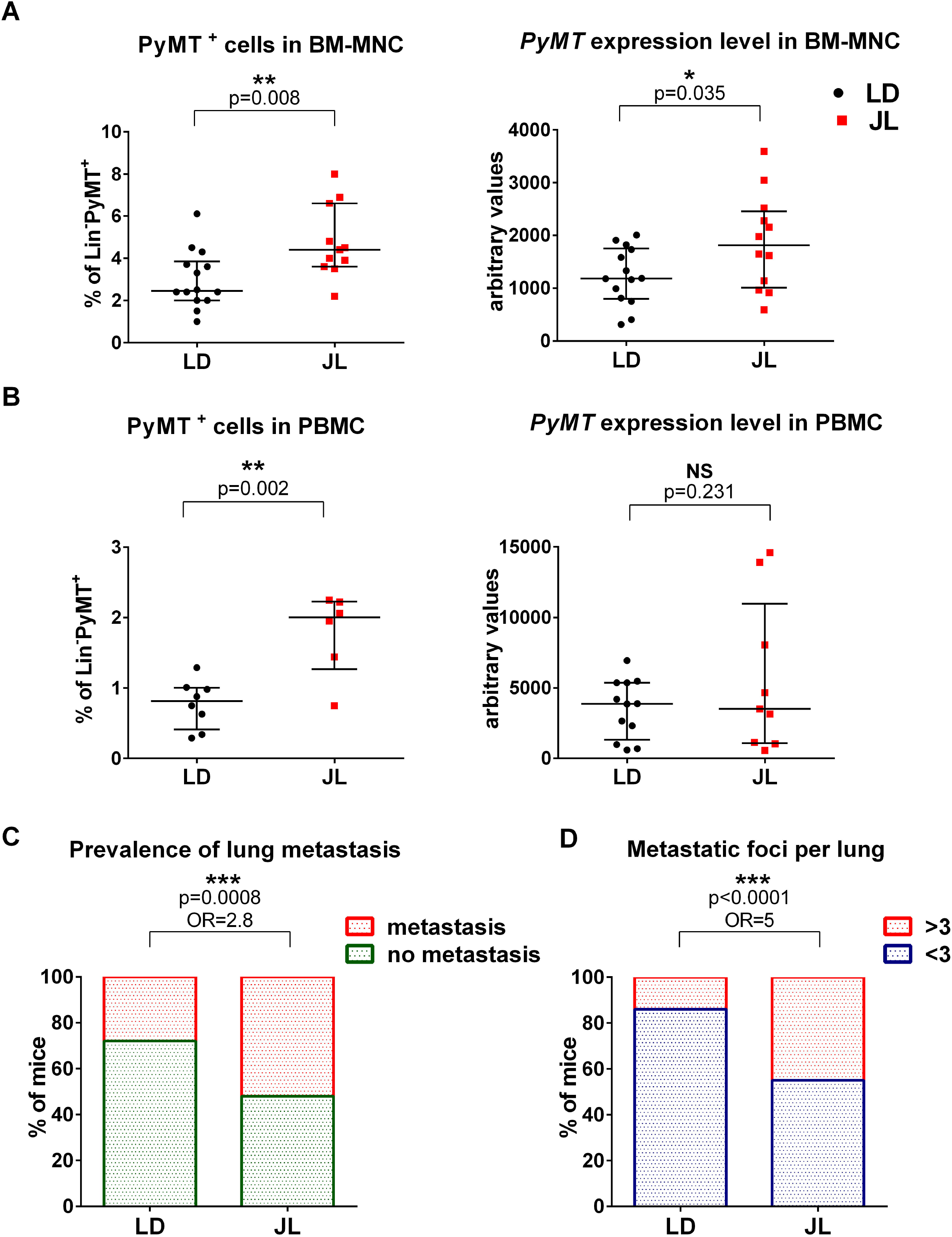
Chronic circadian disruption increases cancer cell dissemination and metastasis formation. (A) Disseminated tumour cells detected in LD (n=14) and JL (n=12) conditions by flow cytometry and real-time PCR in bone marrow mono-nucleated cell compartment (BM-MNC). Data are presented as scatter dot plot with lines representing median with interquartile. p-values are calculated from an unpaired t-test. (B) Circulating tumour cells in peripheral blood PBMCs detected by flow cytometry in LD (n=8) and JL (n=6) and real-time qPCR in LD (n=12) and JL (n=9). Data are presented as scatter dot plot with lines representing median with interquartile. p-values are calculated from an unpaired t-test. (C) Prevalence of lung metastasis in LD (n=25) and JL (n=21) cohorts. To assess prevalence we calculated the percentage of mice with metastasis in LD or JL cohorts. p-value obtained from Chi-square test. Odds ratio presented with 95% confidence interval CI[1.549-5.011] (D) Number of metastatic foci per lung in LD (n=25) and JL (n=21) mice. Data represents the percentage of mice with >3 or <3 metastatic foci in both conditions. p-value obtained from Chi-square test. Odds ratio presented with 95% confidence interval CI[2.524-10.01].

### Gene expression analysis highlights a specific CRD-induced modulation of genes associated with photo-transduction

In order to identify potential molecular players and pathways that drive the increased number of dissemination in JL mice, we performed a mRNA-seq study on bulk of dissociated primary tumour cells and on bulk of BM mononuclear cells from five JL and five LD mice (Supplementary Information, Fig. S4). Hierarchical clustering based on global gene expression profiles clearly separates BM and primary tumours samples but not the LD and JL conditions or the absence/presence of metastasis in lungs (Fig. 3A). We confirmed this observation through principal component analysis on BM and primary cancer cells (Supplementary Information, Fig. S4) where the two first principal components axis do not separate LD and JL samples. Interestingly, in primary tumours, the first principal component axis clearly separates JL mice with or without metastasis (Fig. 3B), indicating a CRD driven mechanism in metastatic spread. We then performed differential gene expression analysis. As expected from the hierarchical clustering, only few genes were differentially expressed significantly between JL and LD conditions (Supplementary Information, Table S2 and Table S3). Intriguingly, in mononuclear BM cells, the most significantly differentially expressed genes (*Rhodopsin, Gnat1, Rbp3* and *Prph2*) are associated with Gene Ontology (GO) terms linked with photoperception and phototransduction (Fig. 3C). They are strongly downregulated in mononuclear BM cells from JL mice, like *Rhodopsin* (23-fold), *Gnat1* (11-fold), *Rbp3* (7-fold) and *Prph2* (6-fold). Similarly, in primary tumours these genes showed significant 3-4-fold downregulation, together with other genes associated with light perception and phototransduction, like *Cngb1, Nrl* (6-fold) or *Bhlhe40* (Fig. 3E). These data suggest a strong effect of CRD on the expression of genes whose function is associated with light perception and light signal transduction, even in tissues that are not supposed to be directly exposed to light (due to poor tissue penetration) or not known to respond to light stimulation.

**Figure 3:**
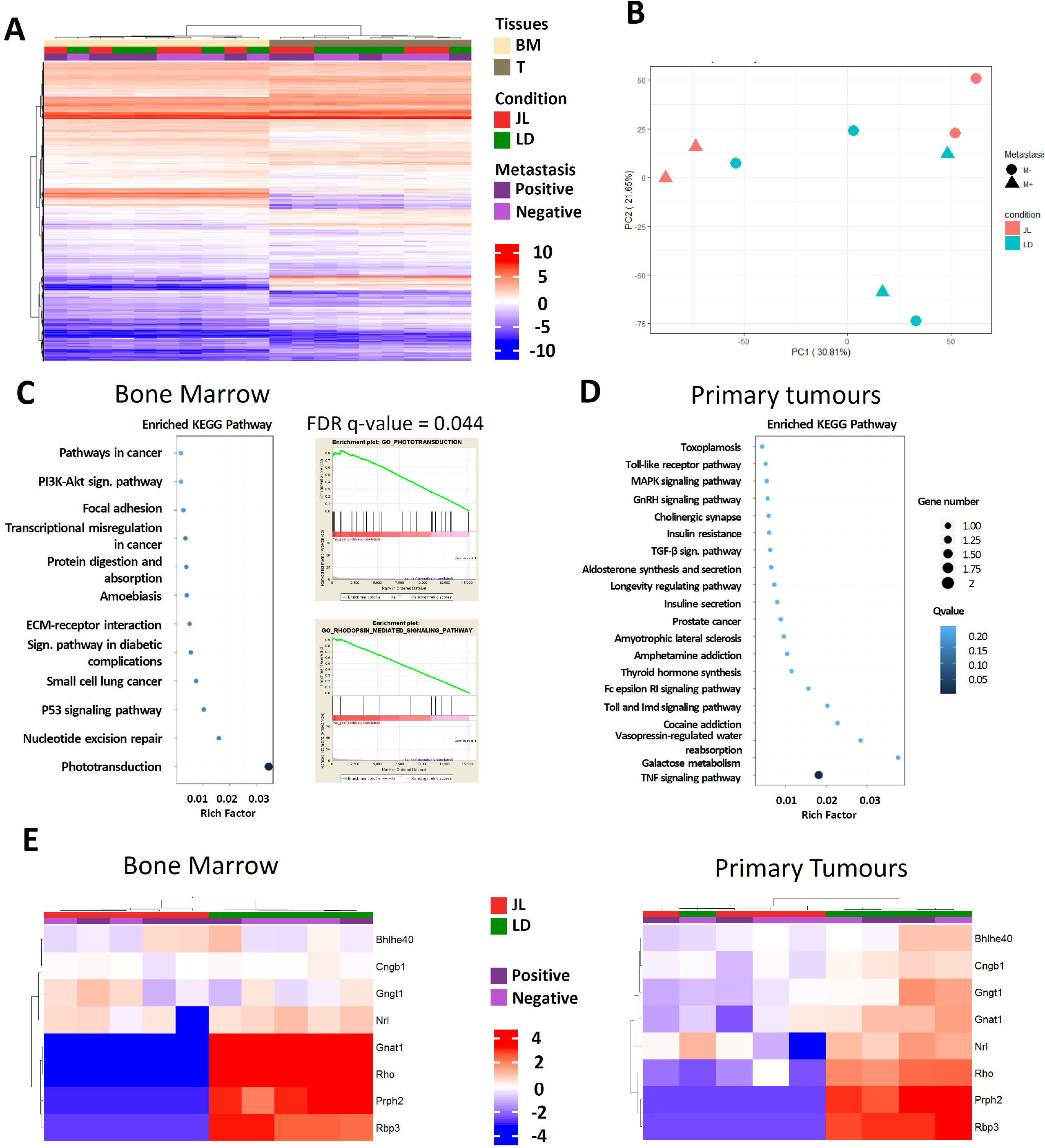
Chronic circadian disruption does not profoundly alter gene expression profiles in primary tumours and Bone marrow mono-nucleated cells except for genes linked with phototransduction and light perception. (A): RNA-seq sample heatmap and hierarchical clustering based on the expression (FPKM) of the 12,556 genes expressed across the two tissues. Gene expression matrix was centered, reduced and log2 transformed. Samples appear as columns and genes as rows and samples are labelled by their tissue of origin (BM: Bone Marrow and T: primary tumour), the experimental conditions (JL in red and LD in green) and the presence of metastasis. Hierarchical clustering is performed using euclidean distance and the Ward.D2 criterion for agglomeration. (B) Principal component analysis (PCA) based on the expression (FPKM) of the 14,487 genes expressed across the nine primary tumours samples. Samples are labelled regarding the experimental conditions (JL in pink and LD in blue) and the presence of metastasis (circle: no metastasis, triangle: positive for metastasis). The percentage of variance explained is indicated for the two first principal components (PC1 and PC2). (C): Enriched KEGG Pathway and Gene Set enrichment analysis for bone marrow. GSEA plots are shown for the GO terms Phototransduction and Rhodopsin_mediated_signaling_pathway. (D) Enriched KEGG Pathway for primary tumours. (E) Heatmap based on the expression of differentially expressed genes linked with phototransduction and photoperception in bone marrow and/or primary tumours. Samples appear as columns and genes as rows and samples are labelled by the experimental conditions (JL in red and LD in green) and the presence of metastasis. Hierarchical clustering is performed using euclidean distance and the Ward.D2 criterion for agglomeration.

### CRD promotes stemness of primary tumour cells

We further characterized the cellular composition of primary tumours in order to understand how chronic CRD can affect the dissemination and metastatic potential of mammary cancer cells. We first tried to assess whether the proportion of mammary cancer stem cells was different between LD and JL conditions. Therefore, we quantified the expression of known mammary stemness markers in cancer cells from primary tumours. Using flow cytometry, we observed a slight increase in the percentage of the cancer cells positive forCD24, CD29, CD49f and CD326 in JL mice (Fig. 4A). In agreement with this data we observed an increase in the expression of *Itgb1* (coding for CD29) and *Itga6* (coding for CD49f) by mRNA-seq (Fig. S5A) together with an upregulation of genes associated with EMT (Supplementary Information, Fig. S5B), a key biological process also known to modulate the stemness of breast cancer cells.^21,22^ The expression of *Inhibin-βA* (*Inhba*) is also increased 4 times in the primary tumours of JL mice. Inhba encodes a protein subunit necessary to to activate TGFβ signalling, a key pathway associated with EMT.^23^ We then evaluated the proportion of the cancer stem cell compartment using a mouse mammary stem cell (MaSC) signature CD49f^hi^CD29^hi^CD24^+/mod^ also described for mouse mammary cancer stem cells.^19,24–26^ We observed a significant enrichment of CD49f^hi^CD29^hi^CD24^+/mod^ cancer cells in primary tumours of JL mice (Fig. 4B). Next, we tested whether this enrichment was associated with an actual increase of stemness potential of primary tumours. To test this hypothesis, we purified cancer cells from primary tumours of LD and JL mice (see methods) and performed mammosphere formation assays. Mammosphere formation efficiency (MFE) was significantly higher in cancer cells from primary tumours of JL mice (Fig. 4C). Previous studies showed that Per2 plays an important role in tumour suppression and that knockdown of *Per2* increases the stemness of mammary epithelial and cancer cells.^13,27^ We also observed a decrease of *Per2* and *Cry2* expression in primary tumours of JL mice (Fig. 4D; Supplementary Information, Table S3) while expression levels of other core clock genes remain similar. We also observed that circadian oscillations of clock genes are modulating the stemness of human mammary epithelial cells. For that, we synchronised MCF12A cells for the circadian rhythm and recovered cells at different times corresponding to *PER2* or *BMAL1* peaks (Supplementary Information, Fig. S5C).^28^ We then assessed the stemness of these cells for the different time points. We observed a diurnal oscillation of mammosphere formation efficiency showing negative correlation with expression peaks of *PER* genes (Fig. 4E). Finally, to quantify the tumour initiation potential of the cells from the primary tumours, we performed orthotopic injection of isolated tumour cells in the mammary fat pad of recipient mice. We observed that the grafted cancer cells purified from JL donor mice show also increased tumour initiating potential in immunocompetent wild type C57BL/6J mice compared to LD (Fig. 4F).

**Figure 4:**
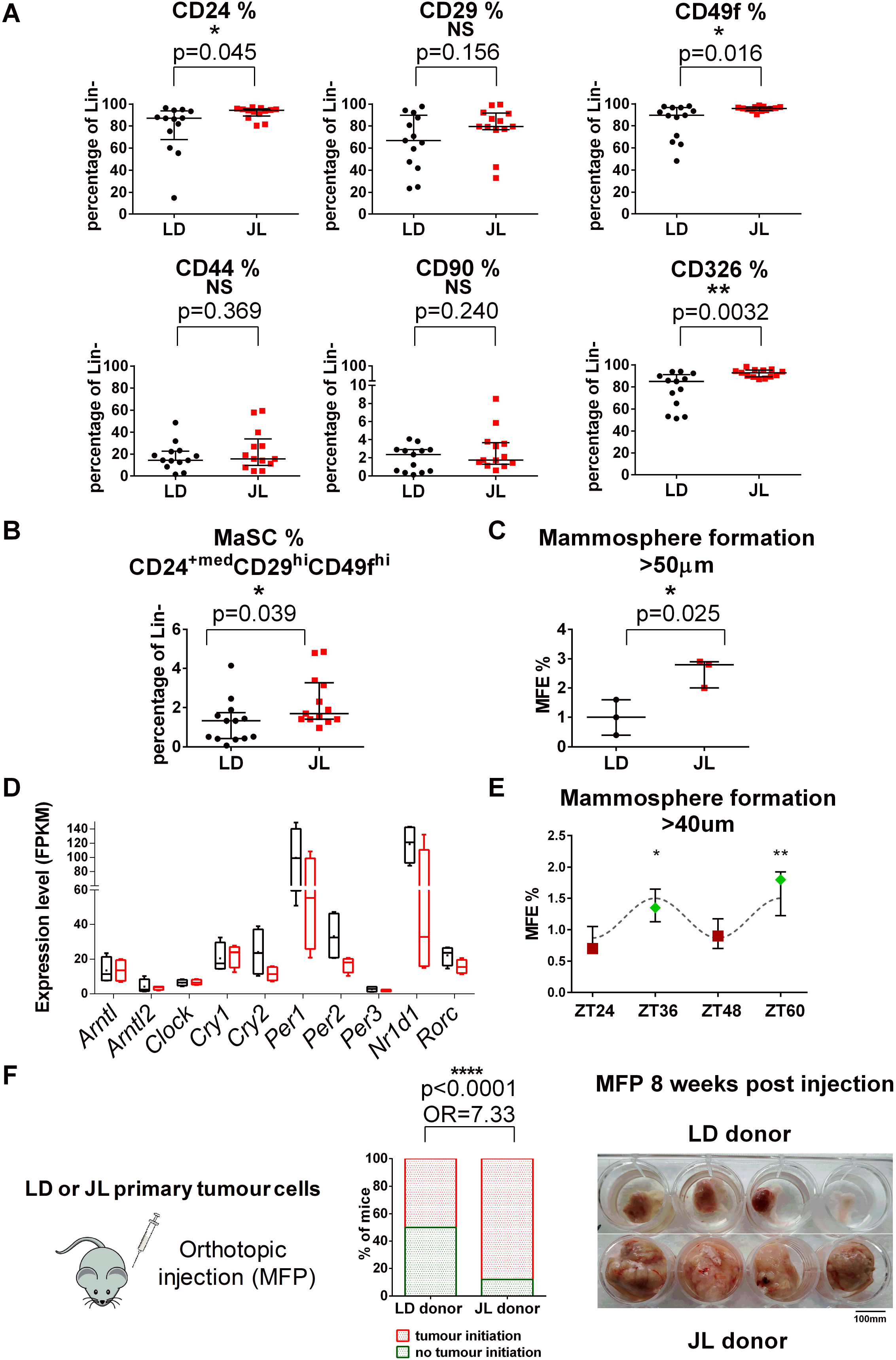
Chronic circadian disruption increases cancer cell stemness in the primary tumours. (A) Relative distribution of stemness marker CD24, CD29, CD49f, CD44, CD90 and CD326 in LD and JL Lin^-^ (CD45^-^CD31^-^CD140a^-^Ter119^-^) tumour cells determined by flow cytometry (n=13). Data are shown as scatter dot plot with lines representing median with interquartile. p-values obtained from an unpaired t-test. (B) Frequency of CD24^+med^CD29^hi^CD49f^hi^ mammary stem cell (MaSC) compartment in LD and JL tumours (n=13). Data are shown as scatter dot plot with lines representing median with interquartile. p-value obtained from an unpaired t-test. (C) Mammosphere formation efficiency (MFE%) of LD and JL tumour cells (n=3). Data are shown as scatter dot plot with lines representing median with interquartile. p-value obtained from an unpaired t-test. (D) Circadian clock genes expression (FPKM) in cancer cells from LD and JL primary tumours. Data presented as box-and-whiskers plots. Variability displayed as medians (line in the box), 25^th^ and 75^th^ percentiles (box) and min to max (whiskers). (E) Mammosphere formation efficiency (MFE%) of MCF12A normal human mammary cell line in different circadian phases (n=3). Red squares and green diamonds represent respectively peaks of *BMAL1*^HIGH^/*PER2*^LOW^ and *BMAL1*^LOW^/*PER2*^HIGH^ expression. Data are presented as dot plot with lines representing median with interquartile. p-value is calculated from one-way ANOVA Tukey’s multiple comparisons test. (F) Tumour initiation study based on orthotopic injection of primary tumour cells from LD (n=6) and JL (n=6) mice in the mammary fat pad of host mice (n=4 per each donor). To assess tumour initiation potential we calculated the percentage of host mice with tumour formation in LD or JL donor cell injected mice. We set a threshold based on MFP weight over 0.25g for positivity. p-value obtained from Chi-square test. Odds ratio presented with 95% confidence interval CI [3.571-15.06]

Altogether these data, in correspondence with previous studies, suggest a regulatory role of PER genes and functional circadian clock in regulating the stemness of mammary epithelial cells that can be altered during CRD.

### Chronic CRD creates an immunosuppressive shift in the tumour microenvironment

The immune system plays a critical role in tumour progression and metastatic dissemination. To investigate whether the increased metastatic prevalence results from changes affecting the tumour immune microenvironment, we characterized the Tumour Infiltrated Cells (TICs) by flow cytometry (see the representative gating strategy on Supplementary Information, Fig. S6). We found reduced number of CD45^+^ immune cells in tumours from JL mice (Fig. 5A) with no significant alteration in the proportional distribution of different immune cell types (Fig. 5B; Supplementary Information, Fig. S7). However, fine characterization of macrophages, dendritic cells (DCs) and T cells sub-populations revealed significant differences between LD and JL tumour immune microenvironment. We identified anti-tumour and pro-tumour macrophage phenotypes based on their MHC II expression level, MHC II^hi^ and MHC II^low^ tumour-associated macrophages (TAMs), respectively.^29^ JL tumours had significantly reduced percentage of the tumour suppressive MHC II^hi,^ TAMs, especially the CD11b^low^MHC II^hi^ TAM1 phenotype (Fig. 5C),^30^ while the tumour supporting MHC II^low^ TAM phenotype significantly increased in these tumours (Fig. 5D). Furthermore, we observed a decrease in the number of infiltrated CD8^+^ T cells and an increase in the CD4/CD8 ratio in JL tumours (Fig. 5E). Significant elevation of CD4/CD8 ratio, which highly correlates with worst prognosis,^31^ was also detected in peripheral blood (Fig. 5F). Our results suggest that chronic CRD weakens the anti-tumour immune response and creates a pro-tumour immune microenvironment. The latter, together with the effects of CRD on cancer cell stemness, may help to facilitate the dissemination of mammary cancer cells and the formation of lung metastasis.

**Figure 5:**
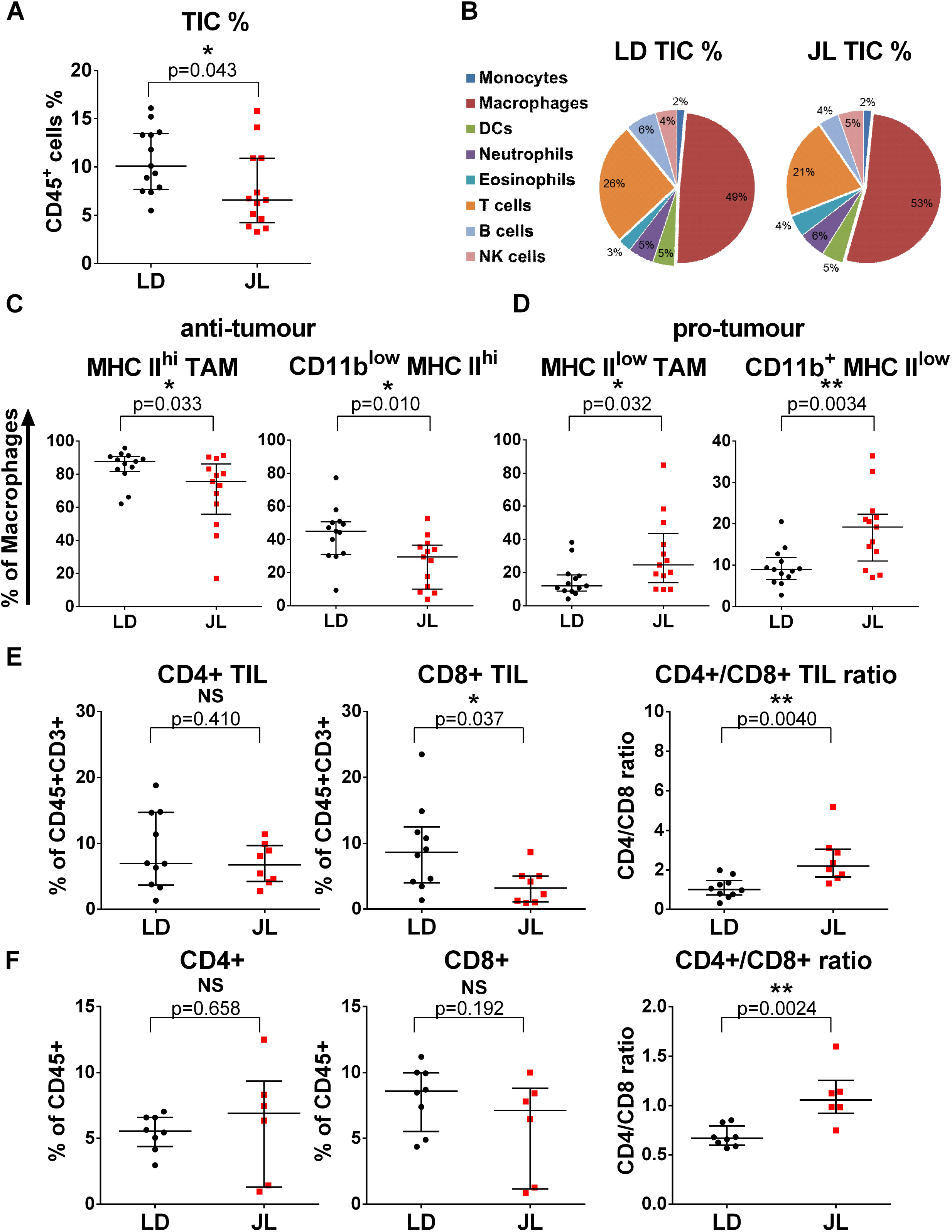
Chronic circadian disruption attenuates immune infiltration and creates a protumour immune microenvironment. (A) Percentage of tumour infiltrated immune cells (TIC) in LD (n=13) and JL (n=13) tumours. Flow cytometric analysis reveals lower percentage of TIC in JL tumours. Data presented as scatter dot plot with lines representing median with interquartile. p-value obtained from an unpaired t-test. (B) Relative distribution of main immune cell types in LD and JL tumours. Data presented as pie charts displaying the mean values of 13 mice. (C-D) Tumour associated macrophage (TAM) phenotypes in LD (n=13) and JL (n=13) tumours: Flow cytometry shows a significant reduction in anti-tumour MHC II^hi^ / CD11b ^low^ MHC II^hi^ phenotype while a significant increase in pro-tumour MHC II^low^ / CD11b^+^MHC II^low^ TAM. Data presented as scatter dot plot with lines representing median with interquartile. p-values obtained from unpaired t-test. (E) Tumour infiltrating lymphocytes (TIL) in LD (n=10) and JL (n=8) tumours: proportion of CD8+ TIL significantly reduced in JL tumours resulting in increased CD4/CD8 ratio. Data presented as scatter dot plot with lines representing median with interquartile. p-values obtained from unpaired t-test. (F) Peripheral blood T cells in LD (n=10) and JL (n=8) mice. Flow cytometry shows an increased CD4/CD8 ratio in peripheral blood of JL mice. Data presented as scatter dot plot with lines representing median with interquartile. p-values obtained from unpaired t-test.

### Chronic CRD alters the cytokine-chemokine network

To better understand how a chronic CRD could modify the tumour immune microenvironment, we characterized the cytokine-chemokine network in JL mice. We first performed a quantification of 17 circulating cytokines using a magnetic Luminex assay on plasma from JL and LD mice. No significant differences were observed, except for IL4 whose circulating levels are slightly decreased in JL mice (Supplementary Information, Table S4). Interestingly circulating levels for CXCL12 were also increased in JL mice even if not significant (p.val=0.2) (Supplementary Information, Table S4). Expression level of cytokines/chemokines and their receptors were also assessed in primary tumours through our transcriptomic study and real-time PCR. We observed that the expression levels of *Ifng, Cxcl13, Tnfs18 or Cxcl11*, known to favour an anti-tumour immune response, were decreased (Fig. 6A; Supplementary Information, Table S3) while the expression levels for *Cxcl3, Cxcl5 and Il1b* were increased (Fig. 6A; Supplementary Information, Fig. S8B, Table S3). As both CXCL12 and CXCL5 together with their respective receptors are described to be involved in metastatic processes ^32–35^ we also assessed the expression level of their receptors, respectively CXCR4 and CXCR2. We observed a significant increase in the number of CXCR4^+^ cancer cells (Fig. 6B) but no differences regarding CXCR2^+^ cancer cells (Supplementary Information, Fig. S8C). Interestingly, we observed a significant increase in the number of CXCR2^+^ TICs in JL tumours (Fig. 6C). These results suggest the existence of an enhanced CXCL5-CXCR2 axis favouring immune cell toward immune suppressive cell types coupled with an enhanced CXCL12-CXCR4 axis favouring cancer cell dissemination. Collectively, these data support the existence of a CRD-driven altered cytokine-chemokine network promoting cancer cell dissemination and immune suppressive tumour phenotype. Therefore to test for CXCR2 effects on tumour progression in JL mice, we used a specific CXCR2 inhibitor (SB265610), ^36,37^ that was injected in MMTV:PyMT mice from 10 to 18 weeks of age and subjected to a chronic jet lag since the age of 6 weeks (Fig. 6D). After the treatment, mice were sacrificed and analysed like previously. We observed a significant decrease of lung metastasis prevalence (Fig. 6E), of PyMT positive DCCs in the BM and blood stream (Fig. 6F) and an increase of TILs (Fig. 6H). T cell infiltration was indeed increased and the CD4/CD8 ratio was decreased indicating an enrichment of cytotoxic CD8 T cells (Fig. 6I). Altogether, treatment with the CXCR2 inhibitor was able to compensate for most of the alterations observed in JL mice regarding cancer cell dissemination and the tumour immune microenvironment.

**Figure 6:**
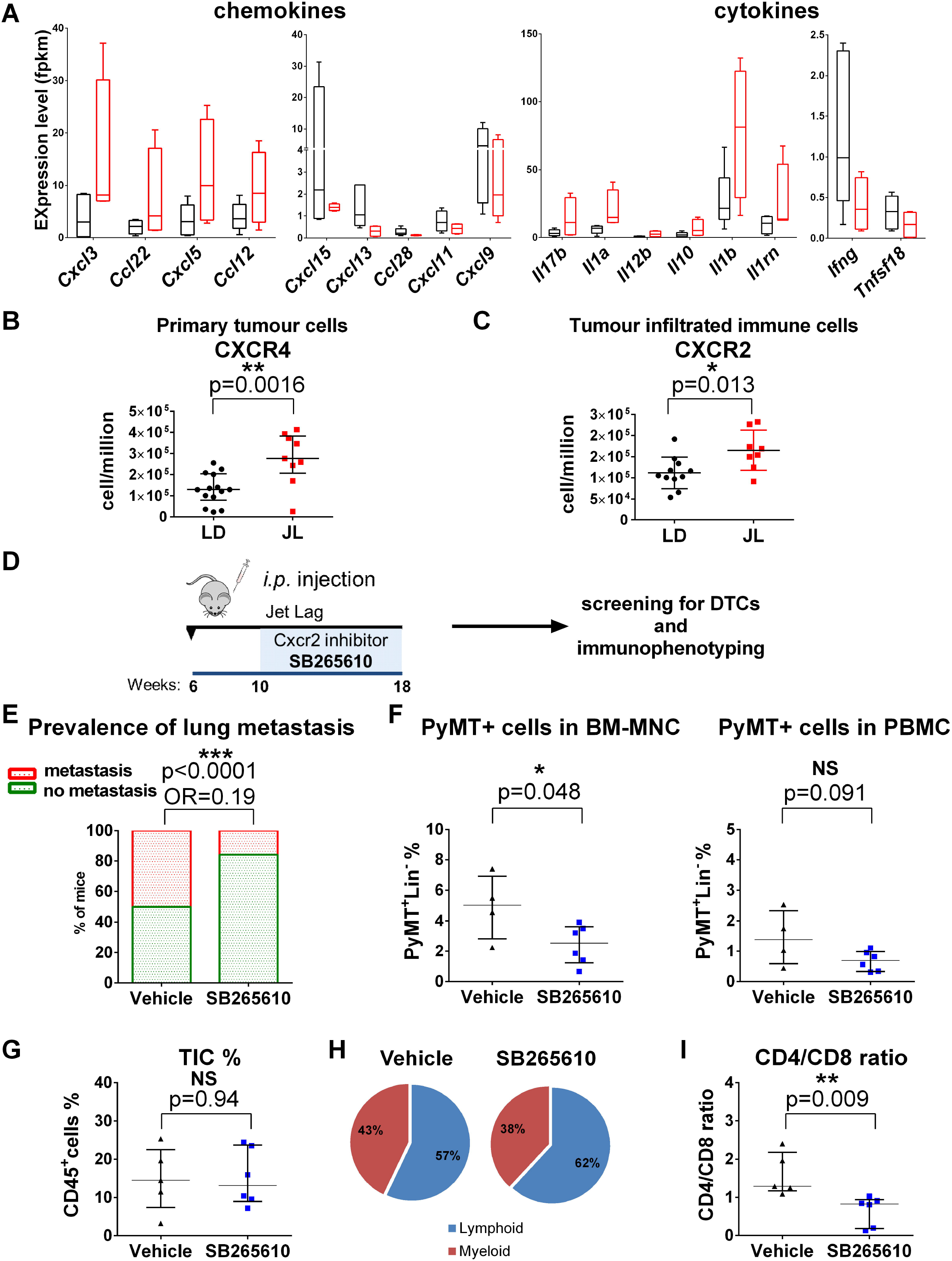
Circadian disruption alters chemokine/cytokine regulatory networks. (A) The most up- and downregulated chemokines and cytokines in LD and JL tumours detected by RNAseq. Expression data represent FPKM values and are presented as box-and-whiskers plot. Variability displayed as medians (line in the box), 25^th^ and 75^th^ percentiles (box) and min to max (whiskers). (B) Number of positive tumour cells positive for CXCR4 in LD (n=14) and JL (n=9) tumours. Data are shown as scatter dot plot with lines representing median with interquartile. p-values obtained from unpaired t-test. (C) Number of positive tumour infiltrated immune cells for CXCR2 in LD (n=11) and JL (n=8) samples. Data are shown as scatter dot plot with lines representing median with interquartile. p-values obtained from unpaired t-test. (D-I) Effects of Cxcr2 axis inhibition on tumour development in JL mice: (D) Scheme illustrating the treatment flow. MMTV:PyMT mice were subjected to a chronic jet lag since the age of 6 weeks and were intraperitoneally (i.p) injected daily (5 days injection + 2 days resting) from 10 to 18 weeks of age with CXCR2 antagonist SB265610 (2mg/kg in 5%DMSO). Mice were sacrificed at age 18 weeks. (E) Prevalence of lung metastasis in mice injected with vehicle (n=5) or SB265610 (n=6). p-value obtained from Chi-square test. Odds ratio presented with 95% confidence interval CI[0.098-0.369]. (F) The percentage of disseminated tumour cells in BM and peripheral blood in vehicle (n=4) and SB265610 (n=6) group. Data are presented as scatter dot plot with lines representing median with interquartile. p-values are calculated from an unpaired t-test. (G) Percentage of tumour infiltrating immune cells (TIC) in vehicle (n=5) and SB265610 (n=6) cohort. Data are presented as scatter dot plot with lines representing median with interquartile. p-values are calculated from an unpaired t-test. (H) Relative distribution of lymphoid and myeloid compartment of TIC in both cohorts. Data presented as pie charts displaying the mean values of 5-6 mice. (I) CD4/CD8 ratio in tumours from vehicle (n=5) and SB265640 (n=6) groups.

## Discussion

In correspondence to epidemiological studies, there is a growing body of experimental evidence linking circadian disruption to increased breast cancer risk and poorer survival outcome.^6,8,38,39^ Here, we present original findings showing that chronic CRD favors cancer cell dissemination and metastasis formation and we shed light on the underlying cellular and molecular alterations. For this purpose we used the MMTV::PyMT spontaneous murine mammary carcinogenesis model,^15,40,41^ which recapitulates many processes involved in human metastatic breast cancer.^20,42^ This model allowed us to work on extended time periods representing a physiological context closer to that of human breast cancer and thus to perform a comprehensive analysis of cancer cells and tumour microenvironment in the view of systematic changes and metastatic dissemination. We applied on this model a jet lag protocol that mimics the effects of shift works or frequent eastbound transmeridian flights and results in severe perturbations in rest-activity, body temperature and clock gene expression in the CNS and peripheral organs.^16^ This protocol also takes into account the observation that the circadian rhythm is more disturbed by advances rather than delays in local time.^43^

Our observations on systemic physiology confirm the importance of circadian rhythm in metabolic processes. In line with previous studies, we found a significant increase of plasma lipid levels in JL mice supporting the link between CRD and cardiovascular diseases.^44–46^ Besides lipid metabolism, feeding-signalling and insulin-glucose axis are also under circadian regulation. Several studies reported elevated leptin and insulin resistance in CRD associated with weight gain, obesity and type-2 diabetes.^14,47–49^ In our study, we detected only fine differences in weight, leptin or insulin levels, which might be due to the timeframe of our study and to the continuous feeding activity of JL mice that reduces physiological differences between rest and activities phases.^50^ Furthermore, we cannot disclose the ability of carcinogenesis to reprogram hepatic homeostasis and metabolism.^51,52^ In this context, we cannot exclude that the developing mammary tumours could partially rewire hepatic circadian homeostasis and consequently lower the CRD induced metabolic changes.

In parallel with the slight metabolic changes observed between conditions, we did not observe major histological differences in primary tumours following 10 weeks of CRD. This supports the fact that the increase in DCCs and the observed enrichment of malignant lesions in JL mice correspond to a global speed-up toward carcinogenesis rather than a selective process leading to the development of different tumour subtypes in LD and JL mice.

Previous studies proposed that CRD boost tumour progression through increased proliferation and metabolic reprogramming.^16,52–55^ Here we demonstrate that CRD also leads to significantly enhanced cancer cell dissemination and metastatic burden. Our results provide experimental evidences that reinforces the findings of previous genetic studies linking cancer severity, relapse and higher risk of metastasis with compromised expression of clock genes.^8,56,57^ We therefore observed reduced expression of *Per2 and Cry2* clock genes in primary tumours of JL mice. Intriguingly, genes involved in phototransduction were among the most significantly altered in both primary tumours and mononuclear BM cells from JL mice. Expression of several phototransduction genes is under circadian regulation through NR1D1 (Rev-Erbα) – NRL, CRX and NR2E3 complexes,^58,59^ and their downregulation is certainly related with CRD, despite the analysed tissues are not directly exposed to light stimuli. In mammals, the functionality of phototransduction molecules in non-visual tissues is poorly investigated. Some data suggests a light-independent activation of phototransduction (transducin/PDE6/Ca2+/cGMP) cascade through Wnt/Frizzled-2 which might function as an anti-apoptotic mechanism.^60,61^ Two recent studies also showed that peripherical clocks can be synchronized by light independently of a functional central circadian clock, and suggested that phototransduction players could drive it.^62,63^

Clock genes and circadian oscillation have been shown to regulate the EMT program,^13,64^ which is one of the mechanisms behind the spread and colonisation of tumour cells ^23,65^. Our differential gene expression analysis of primary tumours revealed an elevated expression of genes associated with EMT in CRD. The upregulated EMT-inducers, *Zeb2, Foxc2* and *Inhba*, have been also implicated in promoting cancer stem cell and metastatic properties.^66–68^ Our data, in consistent with previous studies, support the link between EMT process and stemness/tumour-initiating ability of cancer cells.^21,22,69,70^ We indeed observed enriched stem population in primary tumours from JL mice. Furthermore, our data demonstrate an enhanced stemness and tumour-initiation potential of JL cancer cells *in vitro* and *in vivo*. In addition, we demonstrate here that the stemness of mammary epithelial cells shows diurnal oscillation. It provides further evidence for the regulatory role of circadian rhythm and clock genes on the stemness of mammary cells. The negative correlation between *PER2* expression and mammosphere formation capacity confirms the suppressive effect of Per2 on self-renewal and tumour-initiation.^13^ In line, mammary epithelial cells derived CloctΔ19 mutant mice shows reduced mammosphere formation capacity.^12^ Collectively, these data suggest that CRD promote metastasis by enhancing the EMT program and consequently the tumour-initiating potential of cancer cells.

Circadian regulation of leukocyte homeostasis and immune responses, especially in infections and inflammation, is well documented.^71–73^ However, little is known in the context of tumour microenvironment and tumour immunity. Our results demonstrate that CRD has a profound effect on tumour immunity through modulating the cytokine-chemokine network. Here, we provided evidence that CRD induces a pro-tumorigenic switch of tumour immune microenvironment primarily driven by the altered CXCL5-CXCR2 axis. The circadian expression of CXCL5-CXCR2 axis and its implication in inflammatory diseases have been previously reported.^74–77^ Therefore, in line with previous studies, we propose the following inflammatory cascade as another possible underlying mechanism of CRD related enhanced tumorigenesis and metastatic spread (Fig. 7): CRD increases the expression of Cxcl5 in the tumours leading to enhanced infiltration of CXCR2^+^ myeloid cells, e.g. MDSCs. Consequent accumulation of MDSCs, TAMs and TANs promotes an immunosuppressive microenvironment.^78,79^ These cells are able to directly suppress T cell responses and inhibit CD8 T cell infiltration resulting in impaired anti-tumour activity.^78–82^ Collectively, this autocrine cascade promotes tumour growth and metastasis. In our model, the blockage of CXCR2 enriched the anti-tumour immune cell compartment and significantly reduced metastatic spread, which highly supports our concept. Similarly to our findings, the CXCLs-CXCR2 axis have been implicated in the modulation of tumour immune microenvironment and enhanced metastatic invasion in mouse pancreatic cancer.^34,35^ CXCR2 and several of its ligands (CXCL1, CXCL2, CXCL5, CXCL7 and CXCL8) have been linked to breast cancer progression and metastatic invasion.^83^ It is important to note, that several of human and mouse chemokines are not direct homologues. For example, the murine CXCL5 is most similar to human CXCL5 and CXCL6, however it is also a functional homologue of CXCL8.^84^

**Figure 7.**
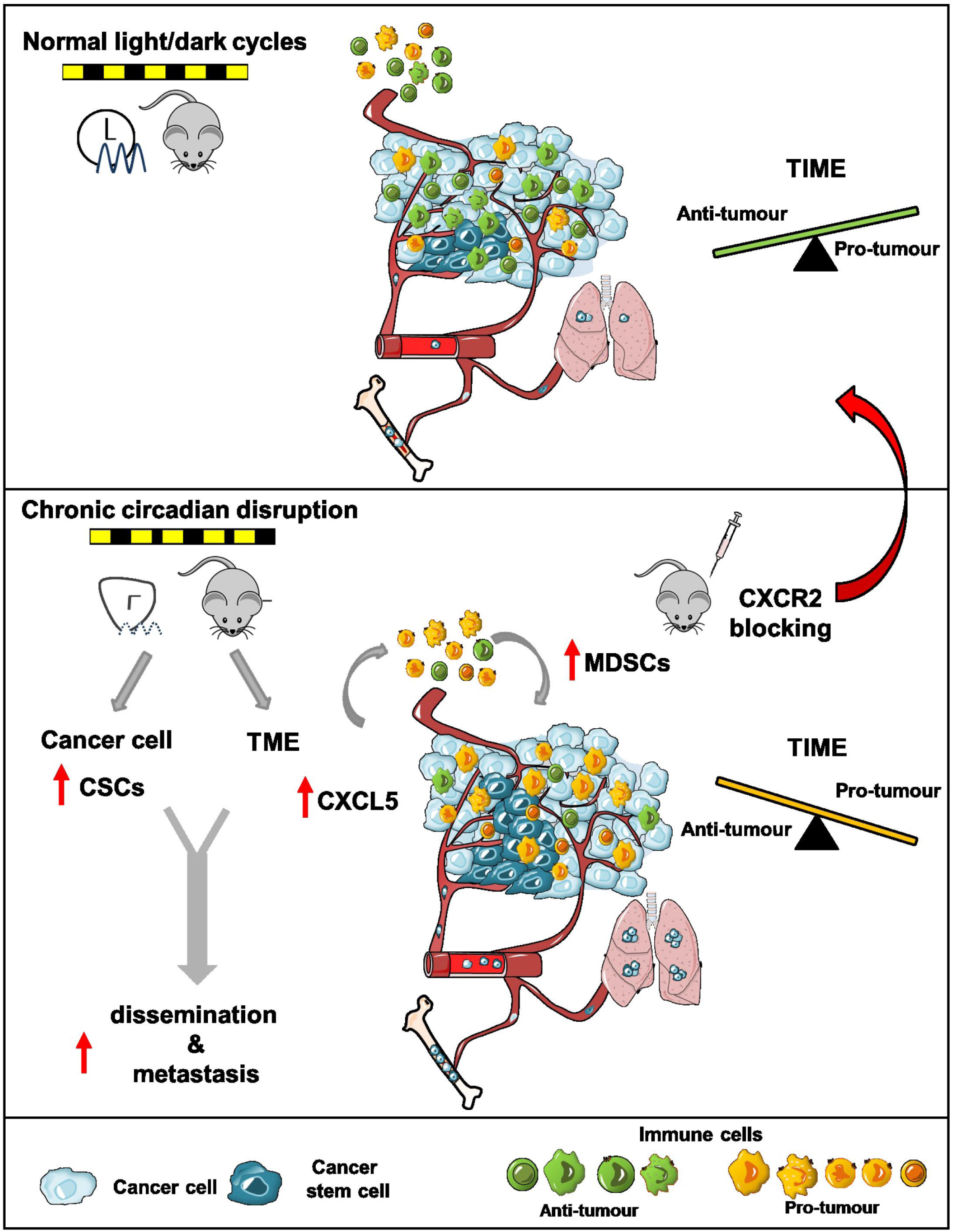
Conceptual schema. Chronic circadian disruption (CRD) alters two key aspects of tumour biology. It increases the proportion of cancer stem cells (CSCs, in dark blue)) and modifies the tumour microenvironment (TME) through the recruitment of myeloid-derived suppressor cells (MDSCs, in yellow) leading to a suppressive tumour immune microenvironment (TIME). This mechanism can be driven at least by an enhanced CXCL5-CXCR2 axis in the TIME. Collectively, these effects result in increased dissemination and metastasis in bone marrow and lungs. Blocking of CXCR2 axis is able to alleviate the effect of CRD and recover anti-tumour activity.

Furthermore, we have also reported an upregulated CXCL12-CXCR4 axis, which is another key mechanism promoting metastatic spread in JL mice. Circadian control of CXCL12-CXCR4 axis^77,85,86^ and its role in immunosuppression and breast cancer metastasis are well described.^32,33,87^

The altered chemokine/chemokine receptor signalling is highly responsible for the CRD induced changes in tumour microenvironment and metastatic capacity. Our findings propose to use CXCR2 inhibition as a novel potential therapeutic tool to thwart CRD effects on tumour progression. Recent studies also suggest the use of CXCR2 or CXCR4 inhibitors in combination with immunotherapy to overcome therapeutic resistance.^82,88^ These novel therapeutic strategies, together with the use of the new predictive tools based on clock genes expression for cancer prognosis will lead to a precision circadian medicine with improved efficacy and immunotherapy responsiveness.^39,89^

In conclusion, our study provides for the first time experimental evidence of the link between CRD and an increase in cancer cell dissemination and metastasis formation. Here we propose two novel molecular mechanisms for how CRD promote tumorigenesis: 1) CRD increases the stemness and tumour-initiating potential of cancer cells, at least by promoting a pro-EMT intratumoral context and 2) CRD reduces anti-tumour immunity through modified chemokine/chemokine receptor signalling. Finally, our findings also point out the potential role of phototransduction players in peripheral tissues impelling further investigation of this topic.

## Methods

### Mice

Mouse strains MMTV-PyMT (JAX #002374) and MMTV-Luc2 (JAX #027827) were purchased from The Jackson Laboratory. Female 5 weeks old C57BL/6JOlaHsd mice were purchased from Envigo (The Neitherlands). Mice were crossed, reared and housed in the institutional animal facility SEIVIL (Service D’Experimentation Animale in vivo INSERM Lavoisier). All mouse experiments were performed in accordance with the DIRECTIVE 2010/63/EU guidelines and were approved by the CEEA26-Paris-Sud Ethics Committee and the French Ministry of Higher Education and Research (APAFIS#8475-2017011015222063 & APAFIS#13183-2018012412194882).

### Cell line culture

The immortalised non-tumorigenic human epithelial cell line MCF12A (ATCC, CRL-10782),^90^ were cultured in DMEM/F12 (Gibco) supplemented with 5% horse serum (Sigma), 10 ng/ml cholera toxin (Sigma), 10 μg/ml insulin (Sigma), 0.5 μg/ml hydrocortisone (Sigma), 20 ng/ml human recombinant epidermal growth factor (StemCell Technologies), 1% Penicillin/Streptomycin (Gibco) and 1% L-Glutamine (Lonza).

### Method Details

#### Jet lag conditions

Mice were kept under normal 12hr light/12hr dark (LD) conditions until 6 weeks age then they were randomly assigned either to remain in LD or to be exposed to jet lag (JL) schedule with 8-hr advance of light/dark cycle every 2 days (Filipski et al 2004, Papagiannakopoulos et al 2016).^16,55^ To induce a chronic circadian rhythm disruption (CRD) mice were kept in JL conditions for 10 weeks in a specific chronobiologic facility and had free access to food and water. Zeitgeber Time (ZT) 0 corresponded to light onset, while ZT12 corresponded to dark onset.

#### Bioluminescence imaging

For *in vivo* imaging of tumours, mice were intraperitoneally (i.p.) injected with 150mg luciferin/kg. Luciferin was resuspended in PBS at 15mg/ml concentration and filter sterilised by 0.22 um filter. Following injection, mice were anesthetised with isoflurane gas (2% during imaging period). Bioluminescence imaging was performed 15, 20, 25 and 30 mins after injection as kinetic analysis determined the peak luciferase activity at 20-25 mins post-injection (IVIS Spectrum imaging system, PerkinElmer). Images were analysed with Living Image software. To determine tumour burden, total flux (photons per second) was measured in a fixed region of interest (whole body).

#### Sample collection

Mice were terminal anesthetised with isoflurane inhalation. Blood was collected by cardiac puncture in EDTA (10% 0.5 M EDTA). Following cervical dislocation tumours/mammary glands, lungs and both hind limb bones were collected and kept in ice cold PBS until further processing.

Blood cell count analysis was performed on 60μl of whole blood using VETSCAN HM5 haematology analyser (ABAXIS). Rest of the blood were span down at 300g for 20 mins at 4□C and 150-300μl plasma were collected and stored at −80□C until further analysis. Erythrocytes were lysed by 1X red blood cell (RBC) lysis buffer (15.5 mM ammonium chloride, 1mM potassium bicarbonate, 10μM EDTA in distilled water). White blood cells were frozen in 10% DMSO/FBS and preserved at −150□C for flow cytometry analysis.

Tumours were weighed to calculate tumour burden (tumour burden % = [tumour weight (g)/body weight (g)] *100). A small piece of tumours was fixed in 4% paraformaldehyde (PFA) for histology. Rest of the tumours were dissociated (see details below).

Lungs were incubated in 1μg/ml heparin in PBS for 1h at 4□C then fixed in 4% PFA for 2-3 days.

Bone marrow (BM) was isolated from femurs and tibias from either both or one hind limbs. BM cells were washed two times with PBS. Erythrocytes were removed by red blood cell lysis. BM cells were frozen in 10% DMSO/FBS and preserved at −150□C for flow cytometry analysis. In cases only one pair of hind limb was flushed out the other were fixed in 4% PFA for histology.

#### Tumour dissociation and cell isolation

Tumours were minced finely with scalpels. Samples were placed into 50ml tube with 10-15ml of pre-warmed dissociation media (1mg/ml collagenase type I, 0.5 mg/ml Dnase I and 1% Penicillin-Streptomycin in DMEM F12) and incubated at 37□C on rotator shaker for 1h. Samples were gently vortexed every 20 mins to avoid clumping. Following incubation, samples were filtered through 70μm nylon strainer and topped up to 40ml with PBS/EDTA (2mM). Cells were span down at 450g for 10 mins. The top fatty layer was collected, topped up to 40ml with PBS/EDTA, gently vortexed and centrifuged. Tumour cells were resuspended in 2ml 1x RBC lysis buffer and incubated for 5 mins to remove red blood cells. Cell suspension was filtered through 40μm nylon strainer. Single cells were frozen in 10% DMSO/FBS and preserved at - 150°C for flow cytometry analysis and tumour initiation study.

#### Detection and quantification of lung metastasis

4% PFA fixed lungs were washed in PBS then permeabilised with 0.1% Triton X/PBS for 2 days. Lungs were incubated in 3% hydrogen peroxide (H_2_O_2_) for 30 mins then the solution was diluted to 1% H_2_O_2_ with PBS and incubated overnight. Blocking was performed with 5% FBS/1% BSA/PBS for 24h. Then lungs were rinsed with PBS then incubated with anti-PyMT antibody (1:100 dilution) for 48h. Following overnight washing, lungs were incubated with rabbit anti-rat IgG AP (1:1000 dilution) secondary antibody for 24h. To reveal antibody labelling lungs were placed into NBT/BCIP working solution. After colour development, lungs were washed and each lobe was imaged by stereomicroscope and metastatic foci were counted. All incubations and washing were done at 4□C.

#### Bone tissue processing and analysis

Tissues samples were fixed in 4% paraformaldehyde overnight at 4 °C. Micro-CT imaging and analyses were performed with a SkyScan 1172 (Bruker microCT), using a voxel size of 12μm, a voltage of 50kV, intensity of 200μA, and exposure of 900ms. NRecon reconstruction and DataViewer software was used for 3D reconstruction. Analysis of structural parameters was performed using CTVox. After micro-CT scan, hind limbs were processed to get histological paraffin sections as follow: Hind limbs were decalcified in 20% EDTA (pH 7.5) at 4°C with constant shaking for 15 days. The EDTA solution was replaced every 2-3 days. Decalcified bones embedded in paraffin were sectioned at 5μm using a Leica microtome. Sections were then stained with hematoxylin and eosin.

#### Flow cytometry

We performed tumour cell characterisation using CD45 PB, CD31 PB, TER-119 PB, CD140A BV421, CD24 BV510, CD29 FITC, CD49f PE, CD44 Pe/Cy7, CD90.1 APC, CD326 APC-Vio770, CXCR1/CD181 AF750, CXCR2/CD182 PE-Vio770 and CXCR4/CD184 APC antibodies. Tumour cells were identified as Lin^-^ (CD45^-^CD31^-^CD140a^-^Ter119^-^) cells. For phenotyping tumour infiltrated immune cells we used CD45 PB, CD24 BV510, CD11b FITC, CD64 PE, CD11c Pe/Cy7, MHC II (IA/AE) APC, Gr1 APC-Vio 770 antibodies and a gating strategy as described on Supplementary Information, Fig. S6. Tumour infiltrating lymphocytes (TILs) were identified using CD45 PE/Dazzle594, CD3 BV510, CD8a APC-Fire750, CD4 FITC antibodies. To perform the immunophenotyping of myeloid cells and lymphocytes from peripheral blood cells and bone marrow cells we used respectively the following combination of antibodies: CD45 PB, Ly6C PerCP/Cy5.5, Ly6G APC-Fire750, CD11b FITC, MHC II APC, CD11c PE/Cy7 and CD45 PE/Dazzle594, CD8a APC-Fire750, CD4 FITC, B220 PE, CD3, CD44 PE/Cy7.

To detect disseminated tumour cells, mononuclear cells from blood or bone marrow were stained for CD45 or CD45, CD31, TER-119 and CD140A respectively. Following cell surface staining, cells were fixed with 1% PFA/PBS for 30mins then permeabilised with 0.1% Triton X/PBS for 20mins. Cells were blocked with 5% FBS/1% BSA/PBS for 30mins. Following PBS wash cells were incubated with anti-PyMT AF488 antibody (1:10 dilution) for 1h. All incubations and staining were done at 4□C.

For live/dead cell separation propidium iodide (PI) or Zombie Violet fixable viability stain were used.

#### RNA isolation and RT-PCR

Total RNA was extracted from bulk of primary tumors dissociated cells and from bulk of bone marrow mononuclear cells using the PureLink RNA kit (Invitrogen) according to the manufacturer’s instructions. cDNA synthesis was performed using High-Capacity cDNA Reverse Transcription kit (Applied Biosystems) with Oligo(dT) primers (ThermoScientific). RT-PCR was performed with QuantStudio 5 (384 well format, Thermo Fisher) using FastStart Universal SYBR Green Master mix (Roche). Samples were used in replicates and melting curve analysis was performed for each run. Geometric mean of *Ctbp1, Prdx1* and *Tbp* Ct values were used for normalisation. Relative fold change (2^-ΔCt^) and gene expression represented as arbitrary units. Primer sequences:

PyMT fw: CTCCAACAGATACACCCGCACATACT, rv: GCTGGTCTTGGTCGCTTTCTGGATAC
*Cxcl5* fw: GGGAAACCATTGTCCCTGA, rv: TCCGATAGTGTGACAGATAGGAAA
*Cxcl3* fw: CAGCCACACTCCAGCCTA, rv: CACAACAGCCCCTGTAGC
*Il1b* fw: GCTTCCTTGTGCAAGTGTCT, rv: GGTGGCATTTCACAGTTGAG
*Ctbp1* fw: GTGCCCTGATGTACCATACCA, rv: GCCAATTCGGACGATGATTCTA
*Prdx1* fw: AATGCAAAAATTGGGTATCCTGC, rv: CGTGGGACACACAAAAGTAAAGT
*Tbp* fw: AGAACAATCCAGACTAGCAGCA, rv: GGGAACTTCACATCACAGCTC

#### mRNA-seq analysis

mRNA-seq was performed by BGI (Hong Kong) using their proper experimental procedures. mRNAs were purified using oligo(dT)-attached magnetic beads. cDNA synthesis was generated from fragmented mRNAs using random hexamer-primers. The synthesized cDNA was subjected to end-repair and 3’adenylation. Adapters were ligated to the ends of these 3’ adenylated cDNA fragments and cDNA fragments were amplified by PCR and purified with Ampure XP Beads (AGENCOURT). Libraries size and quantity were validated on the Agilent 2100 Bioanalyzer. The double stranded PCR product were heat denaturated and circularized by the splint oligo sequence. The single strand circle DNA (ssCirDNA) were formatted as the final library. The libraries were amplified with phi29 to make DNA nanoball (DNB) which had more than 300 copies of one molecule. The DNBs were load into the patterned nanoarray and single end 50 bases reads were generated on a BGISEQ-500 sequencing platform. For each library a minimum of 23M single end reads were produced. All the generated raw sequencing reads were filtered to remove reads with adaptors, reads in which unknown bases are more than 10%, and low quality reads using the SOAPnuke software (https://github.com/BGI-flexlab/SOAPnuke). Clean reads were then obtained and stored as FASTQ format. Reads were mapped using Bowtie2 (Langmead and Salzberg 2012) on the mouse reference genome mm10 (GRCm38), gene expression level was calculated with RSEM ^91^(Li and Dewey 2011) and normalized using the fragments per kilobase of exon per million fragments mapped (FPKM) in each sample. Normalized gene expression values were used to produce heatmaps using the ComplexHeatmap package of R,^92^ using Euclidean distances and agglomeration method Ward.D2 from the hclust function. Principal Component Analysis were performed using pca function of the mixOmics package.^93^ Differentially expressed genes were detected using DESeq2 using un-normalized counts as input.^94^ DEGs were then used to perform pathway functional enrichment with the KEGG annotation database, using the R package hypeR.^95^ List of DEGs were ranked according to p-values and Gene Set Enrichment Analysis (GSEA) were performed using the GSEA software from the Broad Institute.^96^

Raw sequencing data are available under the study accession number: PRJEB33802

#### Immunohistochemistry

4% PFA fixed primary tumours were washed in PBS and transferred to 70% EtOH. They were then embedded in paraffin wax according to standard histological protocols. Five-micrometer thick sections were mounted on adhesive slides (Klinipath-KP-PRINTER ADHESIVES). Paraffin-embedded sections were deparaffinized and stained with HES (hematoxylin, eosin and saffron) to observe the morphology of tumours. All slides were scanned with the Pannoramic Scan 150 (3D Histech) and analyzed with the CaseCenter 2.9 viewer (3D Histech).

#### Tumour initiation study

Cancer cells were enriched by magnetic beads based negative selection using a cocktail of CD45, CD140a, CD31, Ter119 biotinylated antibodies followed by streptavidin microbeads labelling. Flow-through were collected, counted and kept on ice until further use. From each donor 100.000 live tumour cells were injected into the right 4^th^ mammary fat pad of wild type C57BL/6J mice at age 10 weeks. In preparation for injection, tumour cells were resuspended at 5×106 cells/ml concentration in PBS. 20μl of cell suspension was mixed with 20μl of Geltrex (Gibco) and immediately injected into the mammary fat pad using an U-100 insulin syringe. Host mice were kept in LD condition from 6 weeks age until sacrifice at 18 weeks.

#### CXCR2 inhibition

Mice were kept under JL condition, treatment started at age 10 weeks. CXCR2 antagonist SB265610 was resuspended in 5%DMSO-8%Tween80 in 0.9% NaCl and mice were *i.p*. injected daily (5 days injection + 2 days resting) at 2mg/kg concentration. Control group was injected with only vehicle. Mice were sacrificed at age 18 weeks.

#### Plasma chemistry and endocrine analysis

All measurements were performed by the Clinical Chemistry and Haematology platform of Phenomin-ICS (Institut Clinique de la Souris, Strasbourg). Blood chemistry (glucose, albumin, CK, LDH, ASAT, ALAT, ALP, a-amylase, total cholesterol, HDL and LDL cholesterol, triglycerides and creatinine) was performed on an OLYMPUS AU-480 automated laboratory work station (Beckmann Coulter, US) with kits and controls supplied by Beckmann Coulter. Free fatty acid was measured on the AU480 using a kit from Wako (Wako Chemical Inc, Richmond, USA). Internal quality control materials (Olympus) were analyzed on a daily basis to monitor our precision throughout the experiment. Insulin levels were measured on a BioPlex analyser (BioRad) using the Mouse Metabolic Magnetic bead panel kit (Reference: MMHMAG-44K – Milliplex map by Millipore). Corticosterone was measured by RIA using Corticosterone 125I RIA kit for rats & mice (MP biomedical: 07-120102)

#### Luminex assay

Mouse magnetic multiplex Luminex assay for 18 analytes: MCP-1/CCL2, KC/CXCL1, MIP-2/CXCL2, LIX/CXCL5, SDF1/CXCL12, G-CSF, GM-CSF, IFNγ, IL-1β, IL-2, IL-4, IL-6, IL-10, IL-12p70, Leptin, M-CSF, TNFα and VEGF was purchased from R&D Systems-biotechne. Measurement was performed by the Cochin Cytometry and Immunobiology Facility (CYBIO, Institut Cochin, Paris) using Bioplex 200 (Luminex).

#### Cells synchronization for the circadian rhythm

MCF12A cell were entrained by serum shock. Cells were seeded into 60 mm dishes at a density of 200.000 cells /dish). Following 4 days of culture with complete medium the cells were washed with PBS and starved them for 12h in basal DMEM/F1 medium. Thereafter, cells were stimulated with serum shock (complete growth medium supplemented with 50% of horse serum) for 2h. At the end of synchronization (Zeitgeber Time ZT0) cells were washed with pre-warmed PBS and placed in pre-warmed DMEM/F12 supplemented with 20ng/ml hrEGF. Cells were collected at ZT24, ZT36, ZT48 and ZT60 for mammosphere formation assay.

#### Mammosphere formation

For mammosphere formation, we plated either 500 primary murine tumour cells/well or 1000 MCF12A cells/well into ultra-low attachment 24-well plates. Cells were cultured in DMEM/F12 supplimented by 5ug/ml insulin, 4ug/ml heparin, 5ug/ml hydrocortisone and 20ng/ml recombinant murine or human EGF, respectively. Mammosphere formation was performed in 2-4 replicates. Spheres were imaged following 20 days for primary tumour cells and 8 days for MCF12A. Measurement was performed in ImageJ software, threshold was 50μm for primary tumourspheres while 40μm for normal human spheres. Mean values of replicates were used for data analysis. Mammosphere formation efficiency (MFE) was calculated and presented as percentage.

#### Statistical Analysis

GraphPad Prism v6.01 (GraphPad Software, USA) was used to prepare all the graphs and perform statistics, unless stated otherwise. Significance was calculated by t-test and χ^2^ test and one-way ANOVA.

## Data availability

Raw sequencing data are available under the study accession number: PRJEB33802

## Acknowledgements

We thank all the people from the UMRS935 for their technical assistance and daily scientific exchanges. We are grateful to Ibrahim Casals and Benoit Peuteman from the INSERM UMS33 Animal Facility (SEIVIL). We are grateful to Marie-France Champy, Aurélie Auburtin, Tania Sorg and Yann Herault from Phenomin (Institut Clinique de la souris, 1 rue Laurent Fries, 67404 ILLKIRCH cedex 2—CNRS, UMR7104, Illkirch, France—INSERM, U964, Illkirch, France—Université de Strasbourg, France) for their help in blood analysis. We thank to Karine Bailly from the Cochin Cytometry and Immunobiology Facility (CYBIO, Institut Cochin, Paris) for performing the Luminex assay. We thank Francis Levi and Angela Nieto for their helpful comments on the manuscript.

This work was funded by Inserm, University Paris Sud, INRA, Association Institut de Cancérologie et d’Immunogénétique (ICIG), Vaincre le Cancer-NRB, Fond Avenir MASFIP and GEFLUC – Les Entreprises contre le cancer. E.H. post-doctoral fellowship was granted by Vaincre le Cancer-NRB and the University Paris Saclay (Project BioTherAlliance).

## Author contributions

E.H., A.B.G. and H.A. designed the experiments. E.H. performed most of the *in vitro* and *in vivo* experiments with the support of W.T. Y.A. and S.P. performed the analysis of the bone phenotype. M.V., J.L., H.A. and I.R.L. performed the analysis of primary tumour histology. G.D. and C.A. were in charge of animal housing, breeding and manipulation. X.L. and S.D. provided all the protocols, equipment and support for chronobiology experiments. H.A. performed secondary analysis of mRNA-seq data. E.H., S.P. and H.A. wrote the manuscript. A.B.G. and H.A. supervised the research and funded the project.

## Materials & Correspondence

Correspondence and material requests should be adressed to: Hervé Acloque and Eva Hadadi

## Supplementary information

Supplementary Information includes eight figures and four tables.

