## supplementary material for "Chronic circadian disruption modulates breast cancer cell stemness and their immune microenvironment to drive metastasis in mice"

Supplementary information

Supplementary Information includes eight figures and four tables.

**Table S1**: Grades of primary tumours from LD and JL mice according to the PyMT tumour classification by Lin and col. (2003)

**Table S2**: differentially expressed genes in Lin- bone marrow cells between LD and JL conditions.

**Table S3**: differentially expressed genes in cancer cells from primary tumours between LD and JL conditions.

**Table S4**: Cytokines and chemokines quantification using the Luminex technology. UDL: under detection level.

**Figure S1**: Blood biochemistry and hormones levels. Quantification of creatine kinase (CK), creatinine, α-amylase, lactate dehydrogenase (LDH), alanine aminotransferase (ALT), aspartate aminotransferase (AST), AST/ALT ratio, high density lipoprotein (HDL) cholesterol, low density lipoprotein (LDL) cholesterol, adiponectin, leptin, insulin and corticosterone of LD (n=10) and JL (n=10) mice. Data are presented as scatter dot plot with lines representing median and interquartile. p-value calculated from an unpaired t-test.

**Figure S2**: Primary tumour progression was evaluated by routine histology (HES staining). For each experimental condition (LD and JL) sections from four primary tumours from four different animals were analysed and graded according to Lin and col. (2003) in a 4 stage classification scheme that includes hyperplasia, adenoma/ mammary intraepithelial neoplasia (MIN), and early and late carcinoma. For each section, a global view is shown together with two higher magnifications. This highlights the distinct histopathological changes representing morphological events of tumor progression from benign to malignant that are observed in the primary tumor of PyMT model. Scale bars are located on the upper left hand corners.

**Figure S3**: (A) Cancer cell dissemination in bones. (Upper panel) µCT 3D reconstruction (with CTVox program). Lower images are magnification of the green box. Each red arrow indicates a strange whole that could be osteolytic lesions. Each scale represent 1um. (Lower panel) H&E images representing areas, delimited by green dotted lines, containing abnormal cells that can be tumoral cells. BM: bone marrow, DTC: disseminated tumor cells. (B) Quantification of lung metastasis. Lung metastasis were detected using immunocytochemistry and revealed by NBT/BCIP staining (blue labelling, red arrows). Lung metastasis were counted and lungs were classified in three categories: no metastatic foci, between 1-3 and more than 3 metastatic foci.

**Figure S4**: (A) global statistics for reads number and mapping of mRNA-seq data. (B) Principal Component Analysis for all the samples based on common gene expression values (12556 genes). JL samples are in pink and LD in blue. Bone Marrow (BM) samples are represented by circles and primary tumours (T) by triangles. (C) Heatmap based on the expression of all expressed genes associated with the GO term Phototransduction (GO:0007602 ) in mononuclear cells from bone marrow. Samples appear as columns and genes as rows and samples are labelled by the experimental conditions (JL in red and LD in green) and the presence of metastasis. Hierarchical clustering is performed using euclidean distance and the Ward.D2 criterion for agglomeration.

| Mouse number | Experimental conditions (JetLag or LD) | Lung metastasis | number of primary tumours for HES | Grade |
|---|---|---|---|---|
| 1501 | LD | - | 4 | early carcinoma<br>early carcinoma<br>adenoma<br>adenoma |
| 1502 | LD | - | 1 | early carcinoma |
| 1635 | JL | - | 2 | early carcinoma<br>early carcinoma |
| 1636 | JL | + | 3 | early carcinoma<br>early carcinoma<br>adenoma |
| 1639 | JL | - | 1 | adenoma |
| 1643 | JL | + | 2 | late carcinoma<br>late carcinoma |
| 2279 | LD | - | 2 | hyperplasia<br>adenoma |
| 2280 | LD | - | 2 | adenoma<br>adenoma |

### Table S1

| | LD | | JL | | |
|---|---|---|---|---|---|
| | Mean pg/ml | SD | Mean pg/ml | SD | p-value |
| MCP-1/CCL2 | UDL | | UDL | | |
| KC/CXCL1 | 146,6 | 87,38 | 181,5 | 129,1 | 0,5573 |
| MIP-2/CXCL2 | 5,754 | 3,905 | 5,88 | 7,995 | 0,9648 |
| LIX/CXCL5 | 1327 | 668,1 | 1303 | 659,6 | 0,9372 |
| SDF-1/CXCL12 | 255,2 | 140 | 377,9 | 187 | 0,2033 |
| IL-1b | UDL | | UDL | | |
| IL-2 | UDL | | UDL | | |
| **IL-4 **** | 88,44 | 9,23 | 71,52 | 13,24 | 0,0039 |
| IL-6 | 8,112 | 7,581 | 7,272 | 6,295 | 0,845 |
| IL-10 | UDL | | UDL | | |
| IL-12p70 | 115,9 | 75,59 | 171 | 160,7 | 0,4715 |
| IFNγ | UDL | | | | |
| G-CSF | 176,7 | 180,8 | 200 | 168,6 | 0,7699 |
| GM-CSF | UDL | | UDL | | |
| M-CSF | 5,679 | 3,226 | 6,678 | 1,657 | 0,3952 |
| TNFα | UDL | | UDL | | |
| VEGF | 14,77 | 8,201 | 18,14 | 14,09 | 0,5214 |

**Table S4**

Figure S1: blood biochemistry

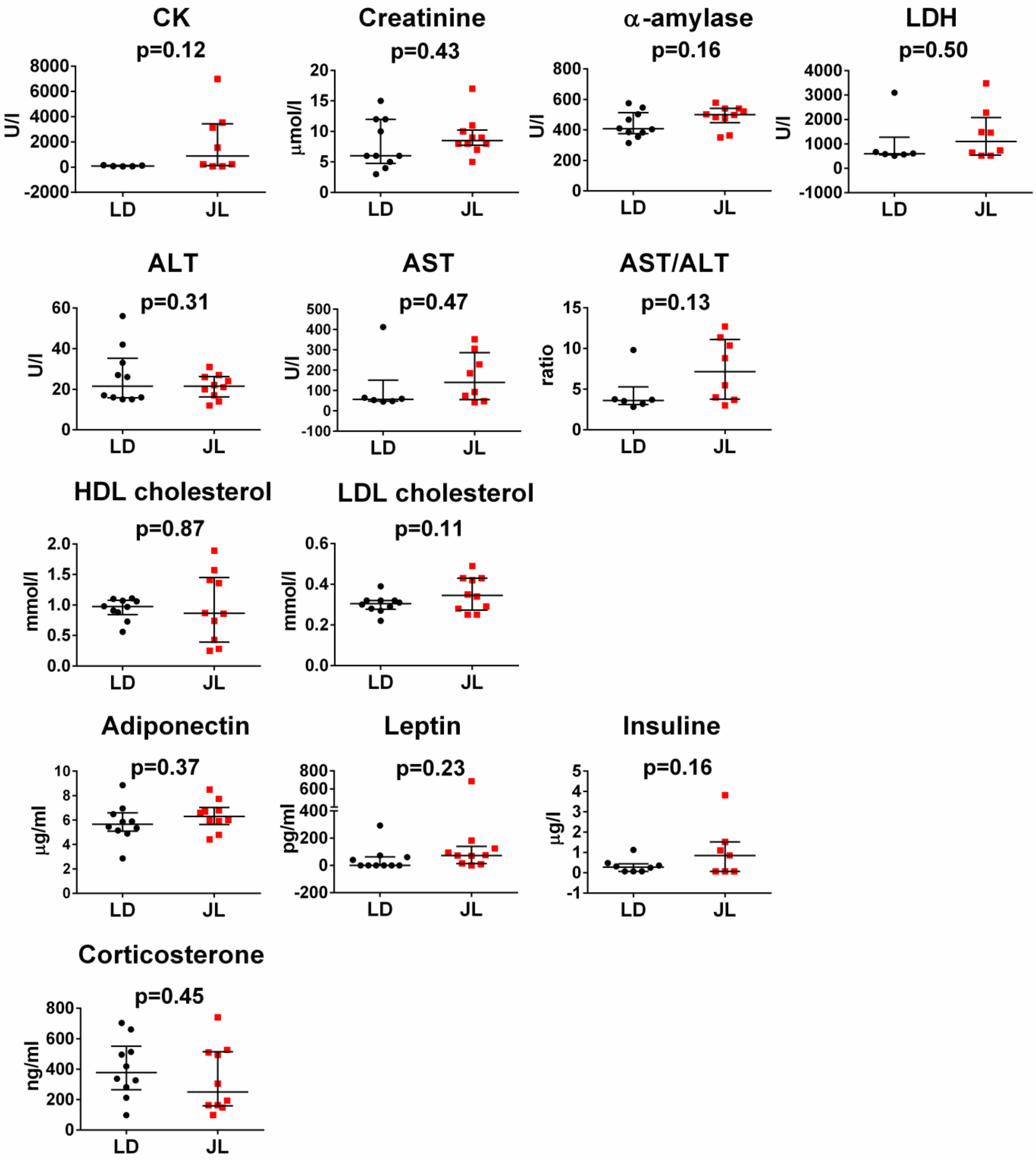

Figure S2: histology (HES staining) of primary tumours

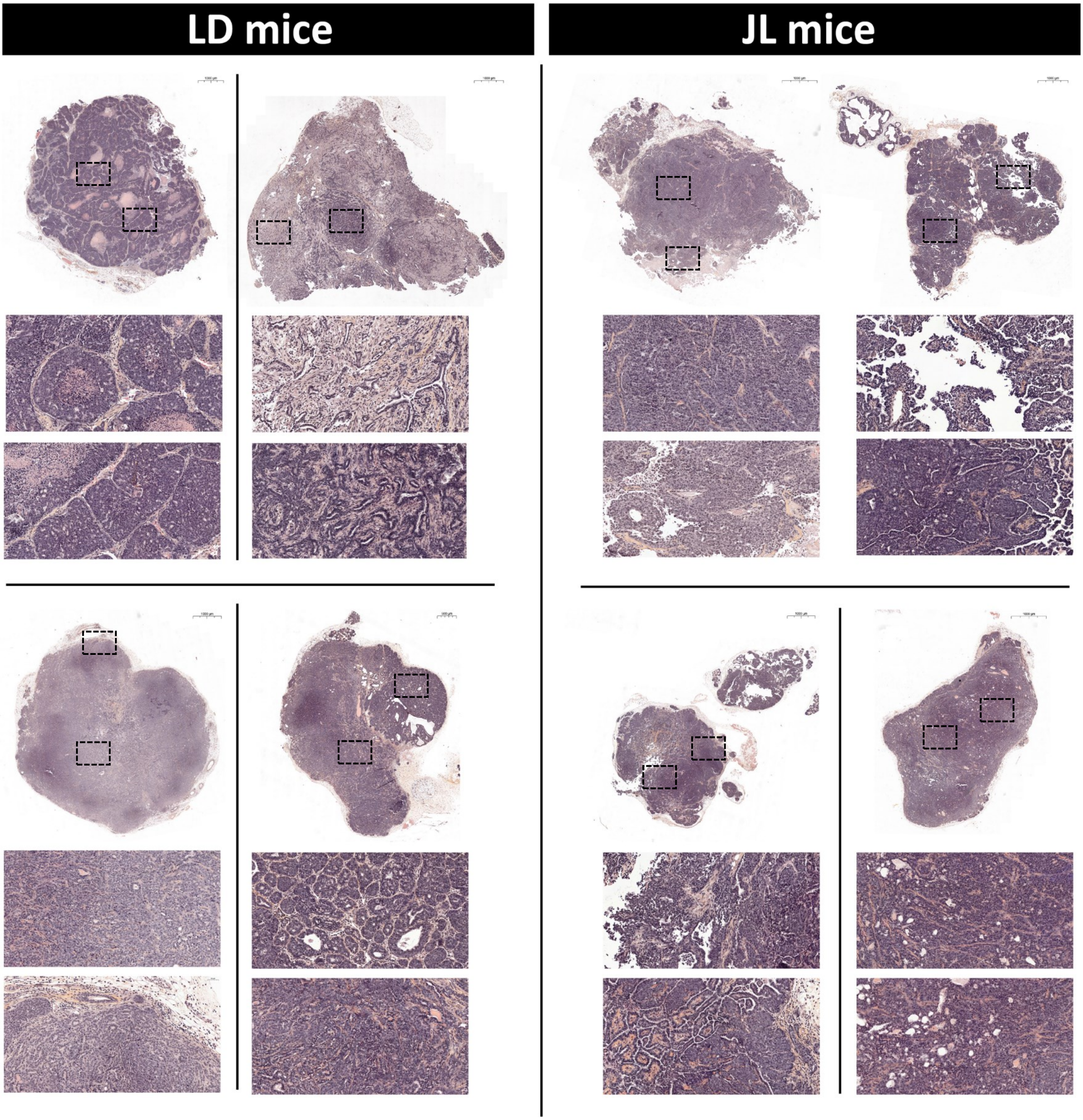

Figure S3: Bone DTCs and Lung Metastasis quantification

**A**

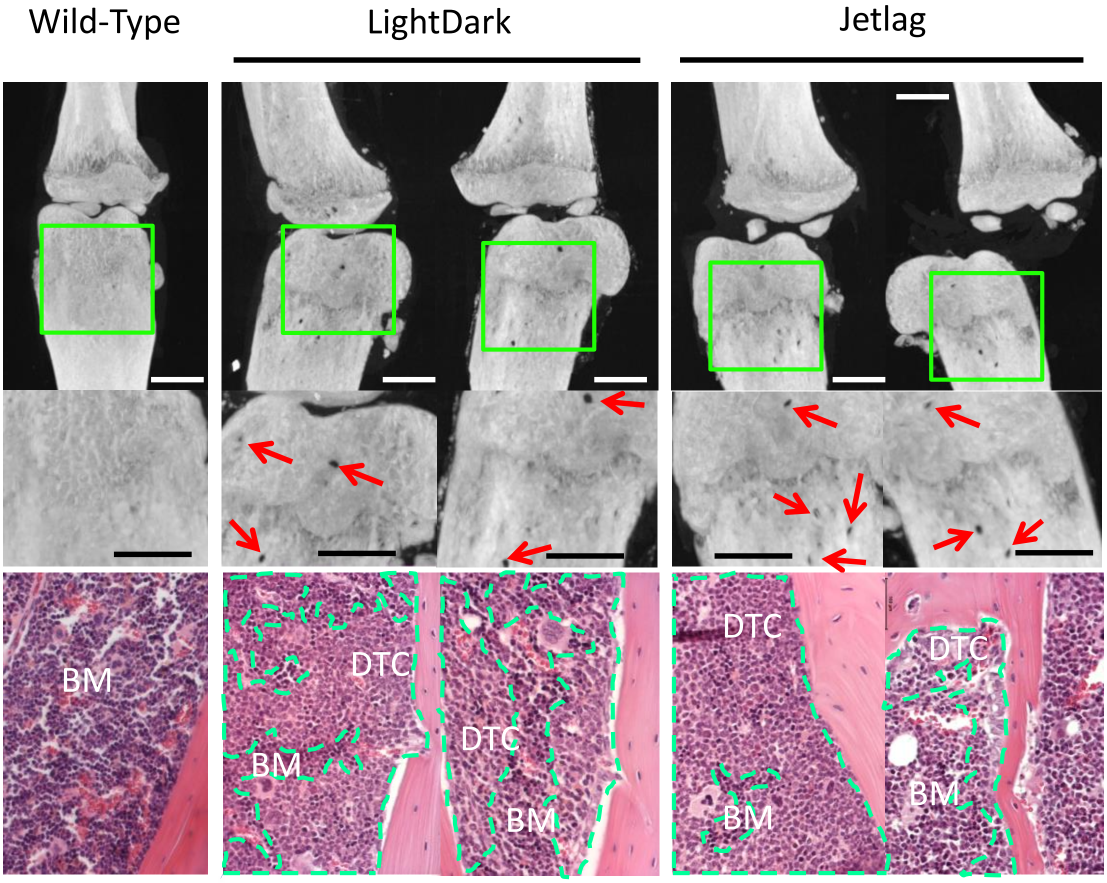


**B**

**Distribution of metastatic foci**

| Conditions | number of metastatic foci | | |
|---|---|---|---|
| | **0** | **1-3** | **>3** |
| **LD** | 18 | 6 | 1 |
| **JL** | 10 | 6 | 5 |

**Macro-metastasis**

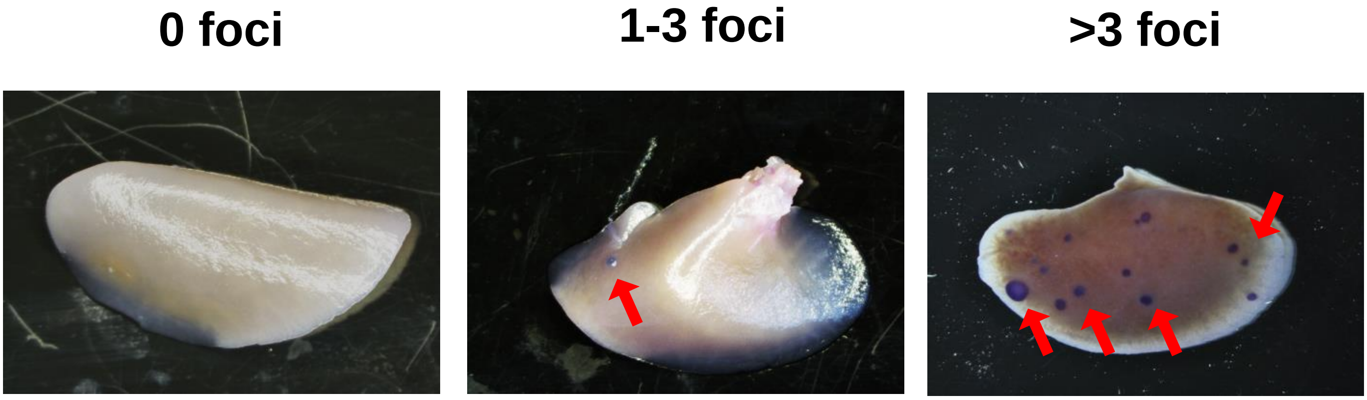

### Figure S4: mRNA-seq analysis

**A**

| Sample | Tissue | Condition | Metastasis | Total CleanReads | Total MappingRatio | Uniquely MappingRatio | Total GeneNumber |
|---|---|---|---|---|---|---|---|
| **1726-BM** | Bone Marrow | LD | M- | 23865426 | 82.97 | 78.68 | 15002 |
| **1726-T** | Primary Tumour | LD | M- | 24124640 | 70.69 | 66.73 | 16137 |
| **1734-BM** | Bone Marrow | LD | M- | 23877046 | 82.68 | 78.46 | 14964 |
| **1734-T** | Primary Tumour | LD | M- | 24071835 | 72.7 | 69.52 | 16253 |
| **1789-BM** | Bone Marrow | JL | M+ | 24097072 | 82.68 | 78.31 | 14481 |
| **1789-T** | Primary Tumour | JL | M+ | 24030875 | 73.61 | 70.2 | 16122 |
| **1791-BM** | Bone Marrow | JL | M+ | 24011010 | 79.74 | 75.55 | 14727 |
| **1791-T** | Primary Tumour | JL | M+ | 24066011 | 63.7 | 60.91 | 15971 |
| **1970-BM** | Bone Marrow | LD | M+ | 24022511 | 82.01 | 77.4 | 15126 |
| **1970-T** | Primary Tumour | LD | M+ | 23939182 | 60.81 | 58.14 | 16373 |
| **2066-BM** | Bone Marrow | LJ | M+ | 24103025 | 81.97 | 77.71 | 15098 |
| **2423-BM** | Bone Marrow | LD | M- | 24070346 | 79.89 | 75.98 | 15391 |
| **2423-T** | Primary Tumour | LD | M- | 24095531 | 55.54 | 53.04 | 16106 |
| **2514-BM** | Bone Marrow | LD | M+ | 24103621 | 83.03 | 78.49 | 15073 |
| **2514-T** | Primary Tumour | LD | M+ | 24125957 | 70.72 | 67.76 | 16517 |
| **2521-BM** | Bone Marrow | JL | M- | 24086082 | 80.86 | 76.58 | 14760 |
| **2521-T** | Primary Tumour | JL | M- | 24048917 | 72.57 | 69.47 | 16434 |
| **2522-BM** | Bone Marrow | JL | M- | 24037282 | 82.8 | 78.48 | 14710 |
| **2522-T** | Primary Tumour | JL | M- | 24078366 | 75.81 | 72.92 | 16315 |

**B**

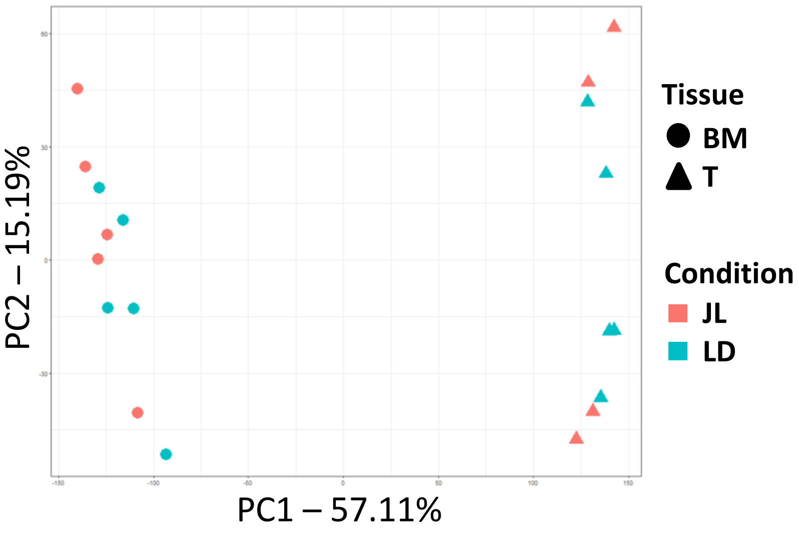


**C** Bone Marrow

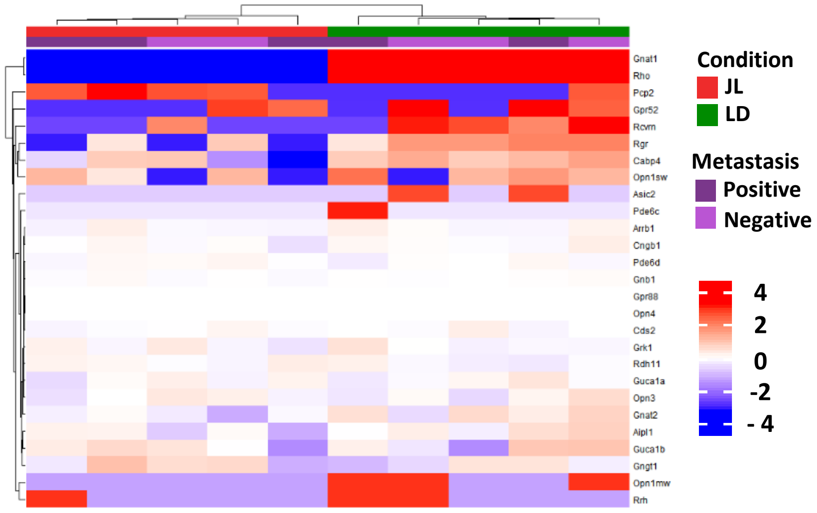

### Figure S5: stemness

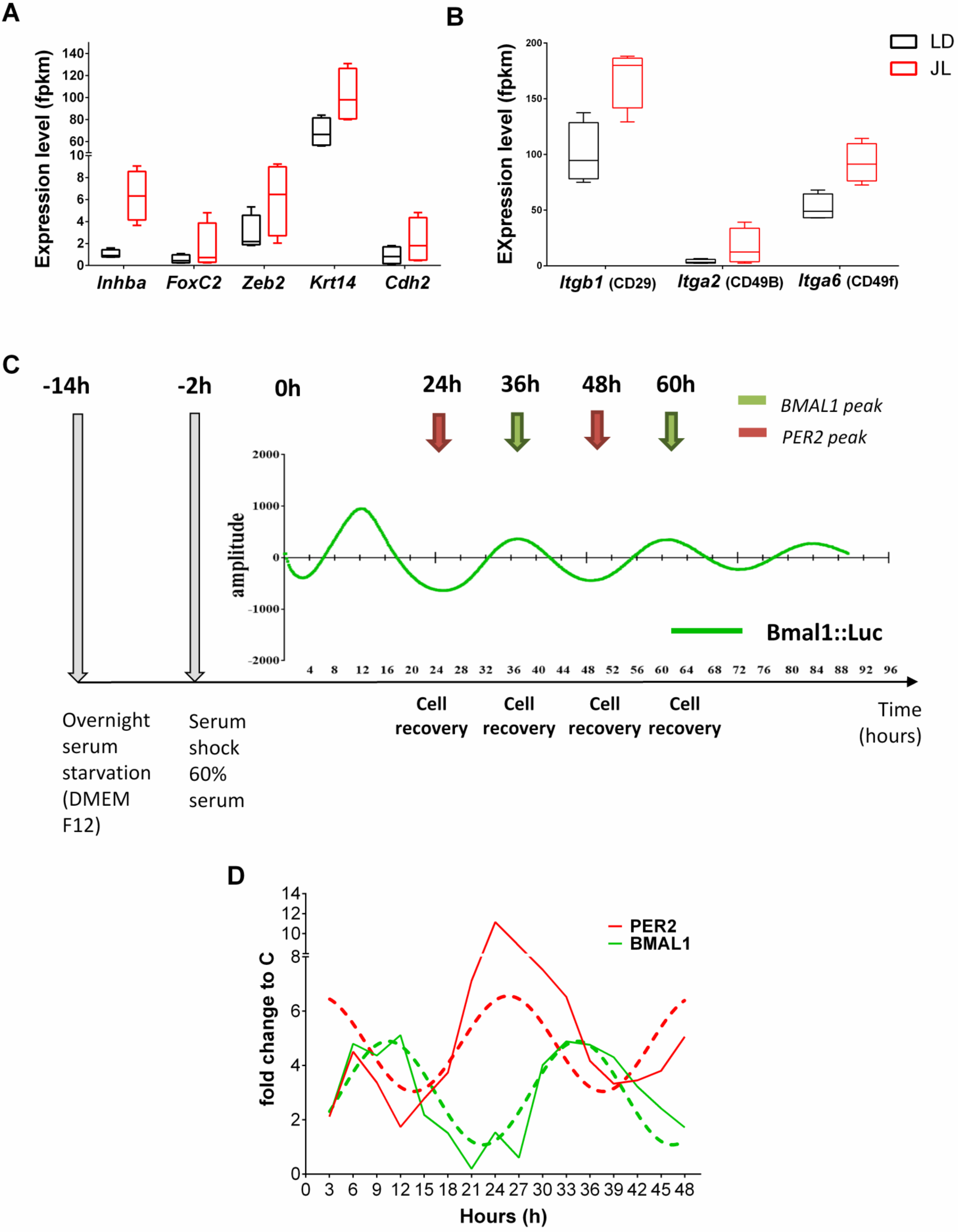

Figure S6: Tumour immunophenotyping

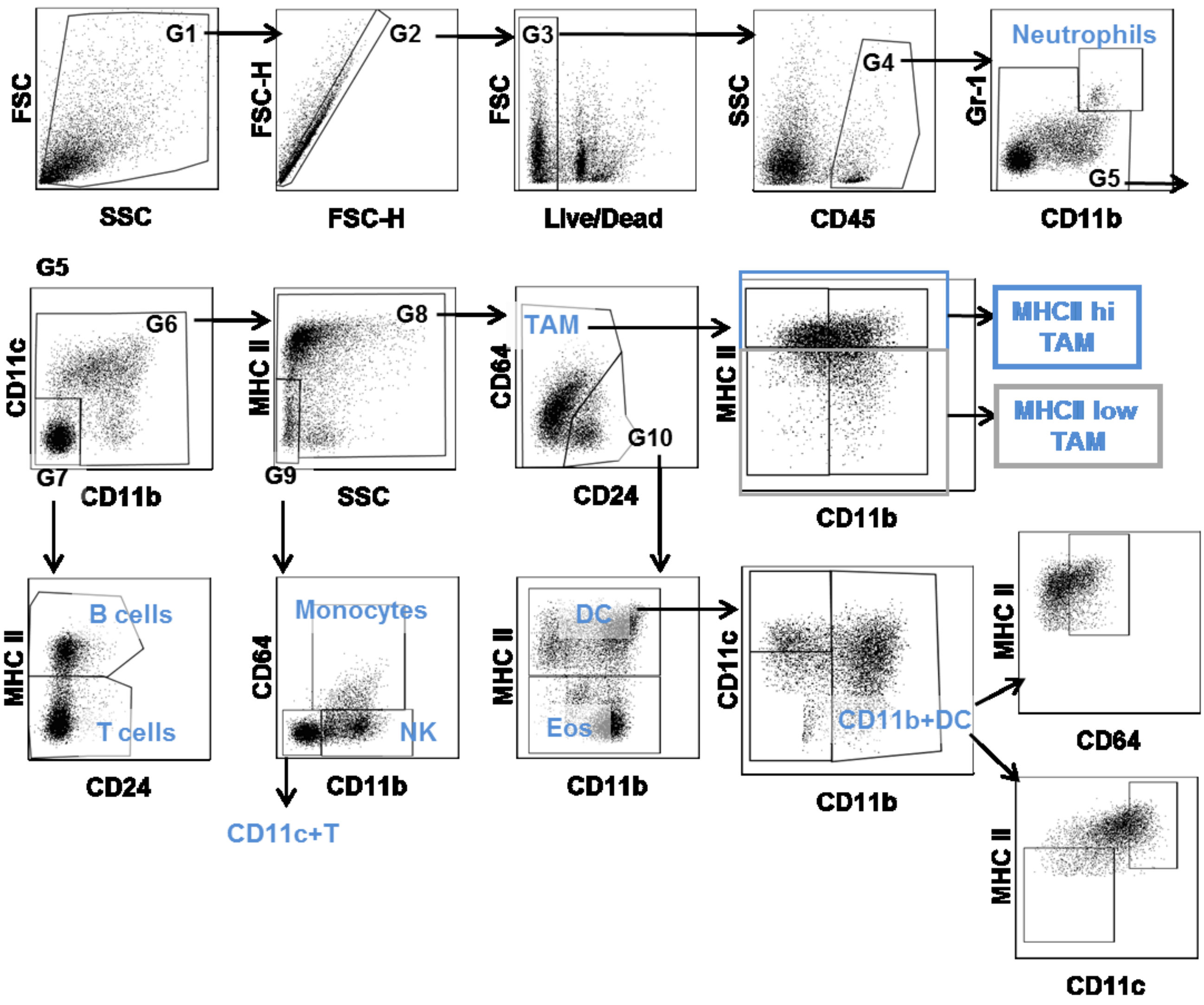

### Figure S7 Main immune cell types in tumours

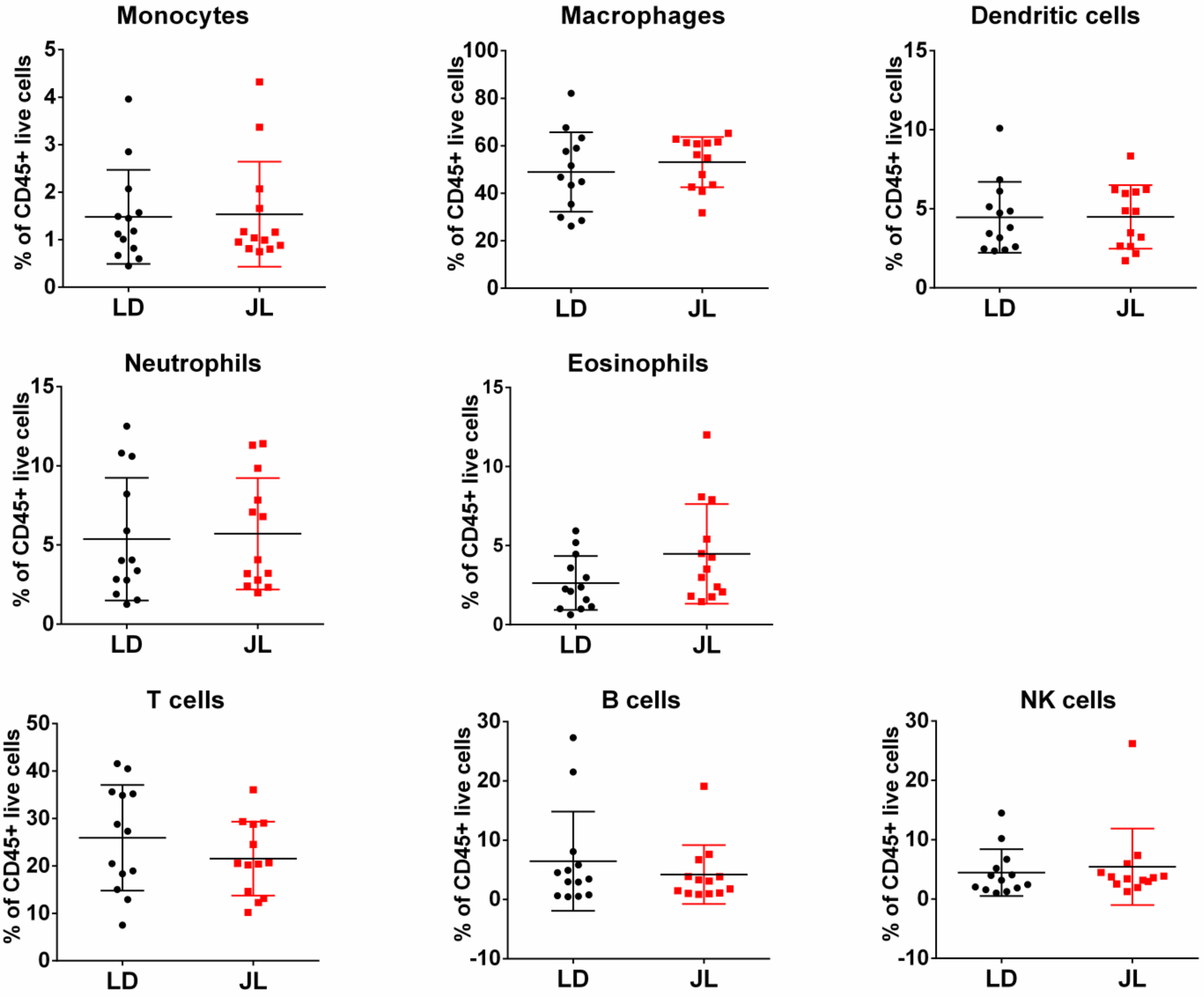

Figure S8: Chronic jetlag alters the cytokine-chemokine network in TME

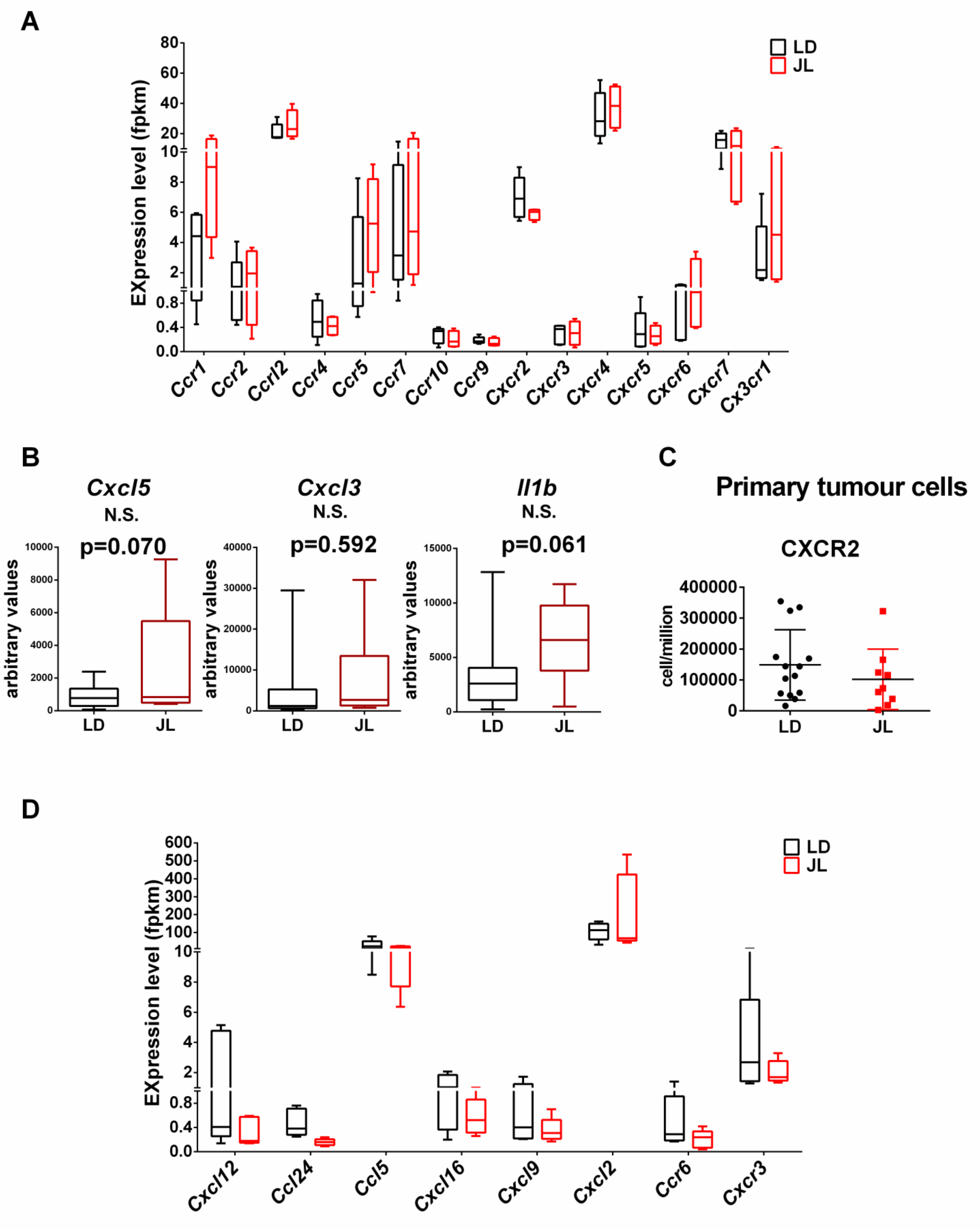

**Figure S5**: (A) Gene expression values (FPKM) of EMT associated genes and of (B) Itgb1, Itga2 and Itga6 in primary tumours from LD (black boxes) and JL (red boxes) mice. Data presented as box-and-whiskers plots. Variability displayed as medians (line in the box), 25th and 75th percentiles (box) and min to max (whiskers). (C) Synchronization protocol used on human mammary epithelial cells to study stemness at different circadian phases. BMAL1 oscillations are represented by the green line using a BMAL1:LUC reporter cell line. Luciferase activity was recorded like in Hadadi and col. (2018). (D) Oscillatory expression profile of PER2 and BMAL1 genes in synchronised MCF12A cells.

**Figure S6**: Gating strategy used for characterizing immune cells

**Figure S7**: Main immune cell types in primary tumours. Proportion of monocytes, macrophages, dentritic cells, neutrophils, eosinophils, T cells, B cells and NK cells were quantified from LD (n=13) and JL (n=13). Data are presented as scatter dot plot with lines representing median and interquartile.

**Figure S8**: (A) Gene expression values (FPKM) of chemokines receptors in primary tumours from LD and JL mice. Data presented as box-and-whiskers plots. Variability displayed as medians (line in the box), 25th and 75th percentiles (box) and min to max (whiskers). (B) Expression values of Cxcl5, Cxcl3 and Il1b quantified by real-time PCR in primary tumour cells from LD and JL mice. Data are presented as scatter dot plot with lines representing median and interquartile. p-value calculated from an unpaired t-test. (C) CXCR2 expression in cancer cells from primary tumours. Data are presented as scatter dot plot with lines representing median and interquartile. (D) Gene expression values (FPKM) of Cxcl12, Ccl24, Ccl5, Cxcl16, Cxcl9, Cxcl2; Ccr6 and Cxcr3. receptors in primary tumours from LD and JL mice. Data presented as box-and-whiskers plots. Variability displayed as medians (line in the box), 25th and 75th percentiles (box) and min to max (whiskers).
